# Characterizing PET CT patterns and bacterial dissemination features of tuberculosis relapse in the macaque model

**DOI:** 10.1101/2025.03.31.646419

**Authors:** Pauline Maiello, Collin Diedrich, Tara Rutledge, Mark Rodgers, Kara Kracinovsky, H. Jacob Borish, Alexander White, Forrest Hopkins, Michael C Chao, Edwin Klein, Sarah Fortune, JoAnne L. Flynn, Philana Ling Lin

**Affiliations:** Department of Microbiology and Molecular Genetics, University of Pittsburgh School of Medicine, Pittsburgh, Pennsylvania, USA; Center for Vaccine Research, University of Pittsburgh School of Medicine, Pittsburgh, Pennsylvania, USA; Department of Pediatrics, Children’s Hospital of Pittsburgh of the University of Pittsburgh Medical Center, University of Pittsburgh School of Medicine Pittsburgh, Pennsylvania, USA; Department of Immunology and Infectious Diseases, Harvard T. H. Chan School of Public Health, Boston, MA, USA; Division of Laboratory Animal Research, University of Pittsburgh School of Medicine, Pittsburgh, Pennsylvania, United States of America; Broad Institute, Harvard University & Massachusetts Institute of Technology; Ragon Institute of MGH, MIT, and Harvard, Cambridge, MA 02139, USA

**Keywords:** Keyword: Tuberculosis, Relapse, PET CT, HIV

## Abstract

Tuberculosis (TB) relapse after appropriate drug treatment is poorly understood but critical to developing shorter treatment regimens. Using a cynomolgus macaque model of human TB, macaques with active TB disease were treated with a short course of isoniazid and rifampin and subsequently infected with SIV. Serial clinical, microbiologic, immunologic and position emission and computed tomography (PET CT) assessments were performed to identify risk factors of relapse. Of the 12 animals, eight developed radiologically defined relapse including four that had clinical and/or microbiologic signs. Greater gross pathology and bacterial burden were observed in relapse animals. PET CT characteristics before, during, and at the end of treatment were similar amongst relapse and non-relapse animals. We show that complete sterilization or very low Mtb burden is protective against SIV-induced TB relapse but cannot be predicted by PET CT. Using bar-coded *M. tuberculosis*, we found that Mtb dissemination during relapse originated from both lung and thoracic lymph nodes, underscoring the importance of lymph nodes as a reservoir. By matching bar-coded Mtb and serial PET CT, we also demonstrate that not every site of persistent Mtb growth after drug treatment is capable of dissemination and relapse, underscoring the complex nature of drug treatment and relapse.

## Introduction

In 2023, ∼8.2 million people were newly diagnosed with symptomatic, active tuberculosis (TB) caused by *Mycobacterium tuberculosis* (Mtb), and 1.2 million deaths occurred(1). Standard TB treatment involves 6 months of multiple drugs and has a cure rate of 85%(1). But an estimated 7% of all patients develop TB relapse(2), though relapse from primary infection after treatment can be difficult to distinguish from reinfection without molecular testing of serial samples(3). In meta-analysis from 1980-2020 addressing reinfection/relapse within 2 years, 70% of cases were from relapse and 30% from reinfection(4). Using a multi-predictor analysis from large pool of published patients, HIV infection (OR, 2.6,95% CI 1.4-4.6), baseline cavitary disease with a positive smear at 2 months of treatment (OR 2.3,95% CI 1.3-4.3), and baseline cavitary disease with a positive culture at 2 months post treatment (OR 2.1, 95% CI 1.2-3.8) had the highest odds of relapse, but these factors only account 10% of all relapse cases(2). TB relapse is clearly underappreciated and not well understood.

Given that the majority of relapse cases have no identified risk factor, better models and tools are needed to understand and prevent relapse (reviewed in(5–7)). Positron emission tomography (PET) using ^18^F-fluorodeoxyglucose (FDG) as a metabolic PET probe with computed tomography (CT) has been used to follow TB disease progression/treatment with promising results in predicting treatment success(8–11). This is particularly important as the development of shorter treatment regimens is a key TB eradication strategy, but long term patient follow up is difficult and expensive. Models that can predict TB cure and improve our basic understanding of the events that control and prevent relapse are direly needed.

Serial FDG-PET CT in the macaque model of TB can be extremely useful in understanding human Mtb infection, HIV-TB coinfection, response to drugs, and vaccine efficacy(12). Recognizing that HIV is a major risk factor for relapse, we modified our TB-SIV coinfection animal model with the aim to examine TB relapse. Cynomolgus macaques infected with low dose Mtb develop the full spectrum of outcomes and pathology seen in humans(13–15). Macaques with clinically defined active TB were treated with isoniazid (INH) and rifampin (RIF) for two months followed by a one month wash out period and then SIV infection while serial assessments were performed. Eight of the 12 animals developed radiographically defined relapse resulting in greater gross pathology disease and bacterial burden. Using bar-coded Mtb and PET CT, we provide insights into the dynamic and complex nature of bacterial dissemination during relapse.

## Methods

### Animals and study design

Adult Chinese cynomolgus macaques (*Macacca fasicularis*) (Valley Biosystems, Sacramento, CA) were screened for other infectious co-morbidities (e.g. SIV, SRV, parasites, Mtb) before infection. Animals were infected with molecularly bar-coded (∼15 CFU) Mtb Erdman strain via bronchoscopic instillation that was confirmed by PET CT scan(16). Animals with clinically defined active TB disease(13, 15) (i.e., having signs of disease [cough, weight loss, anorexia], evidence of Mtb growth by gastric aspirate [GA] or bronchoalveolar lavage [BAL], and progressive disease by PET CT imaging) were treated with human equivalent doses of isoniazid (INH, 15 mg/kg/dose daily by mouth) and rifampin (RIF, 20 mg/kg/dose daily by mouth) for 8 weeks. All animals received at least 85-100% of their doses. After a 1 month drug-free period, animals underwent SIV_mac251_ infection (swarm) (1.67×10^5^ viral RNA copies, intravenous) for 8 weeks until necropsy (Figure 1A). (See Supplementary Table 1.) Serial clinical, immunologic, microbiologic, and radiographic assessment were performed for the entire duration of the study. Blood was obtained via venipuncture for isolation of peripheral blood mononuclear cells (PBMC) every 1-4 weeks and bronchoalveolar lavage (BAL) monthly for flow cytometric analysis as previously described(17).

**Figure 1.**
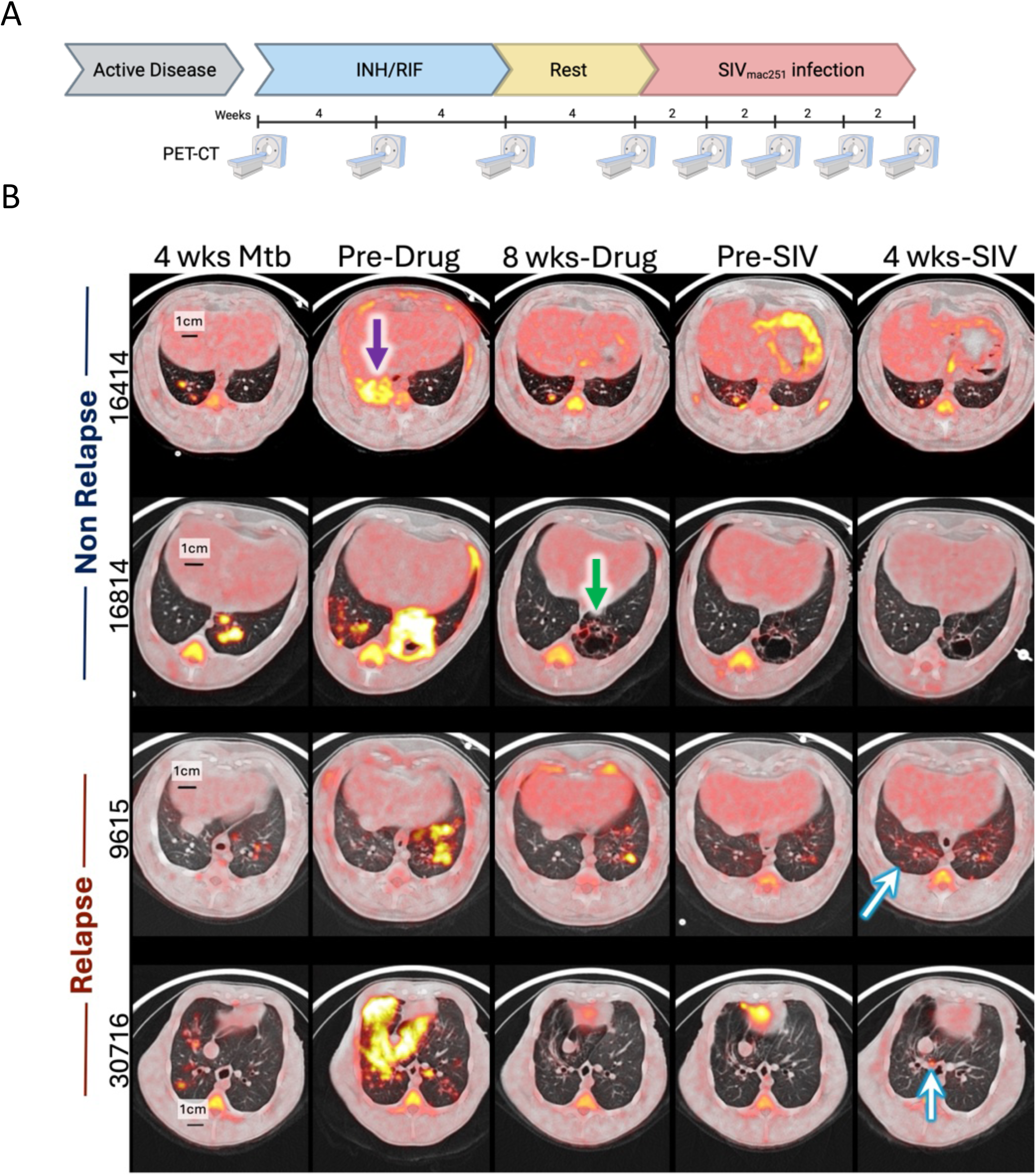
Experimental design and serial PET CT images from animals with active TB before and during treatment and relapse. (A) Experimental study design. Animals with active TB disease were given a 2 months course of isoniazid (INH) and rifampin (RIF) followed by 1 month of no drugs after which SIV infection occurred. Serial PET CT was performed prior to, during drug treatment, before and after SIV infection as shown above. Created with Biorender.com (B) Serial PET CT images of TB disease in cynomolgus macaques at 4 weeks post-Mtb infection, pre drug treatment, 8 weeks post-drug treatment, pre-SIV infection, and 4 weeks post-SIV infection. Axial images of the chest demonstrate the degree of TB-associated lung involvement and FDG activity (shown in yellow) among 2 animals without evidence of relapse (16414 and 16814) and animals with relapse (9615 and 30716). Purple arrow shows a consolidation and green arrow shows a cavity. Blue and white arrows point to granulomas that appeared on scan after SIV infection.

### PET CT acquisition and analysis

*In vivo* progression of disease was monitored and quantified using the microPET Focus 220 preclinical PET scanner (Siemens Medical Solutions) and a clinical 8 slice helical CT scanner (NeuroLogica Corp) with previously published methods and metrics(18). Scans were performed before treatment and every 2-4 weeks thereafter. (No scans were performed prior to Mtb infection due to budgetary constraints.) PET CT analysts (PM, AGW, HJB) were blinded to treatment group. Relapse was defined as the development of new lung lesions identified by PET CT (and confirmed at necropsy) after completing INH/RIF treatment.

### Quantification of Disease, Mtb burden, SIV RNA, and Mtb genome isolation with bar-code mapping

Gross pathology of TB disease was quantified at necropsy using a previously validated method (19). Scan matched lung granulomas and other pathologies from other sites (e.g., lymph nodes, liver) were harvested. Tissues were homogenized into single cell suspension for immunologic assays and bacterial burden(19). Mtb burden was quantified in colony forming units (CFU) from individual sites(19). Lymph node burden (“LN CFU”) is the sum of CFU from all thoracic lymph nodes. Extrapulmonary score (EP score) is a quantitative estimate of extrapulmonary involvement (e.g., liver, paracostal abscess, kidney) based on bacterial growth and gross or microscopic evidence of TB(19). Total bacterial burden includes the sum of CFU from the lymph nodes (thoracic and extrapulmonary) and thoracic cavity (grossly normal lung, granulomas, involved lung, or diaphragm granulomas). Bacterial burden for animal 16714 was omitted due to contamination. Samples with Mtb growth were sequenced for bar-code identification using prior methods(16). Each bar-code from disease sites was mapped three dimensionally (estimated Cartesian coordinates by PET CT) using the Osirix imaging software (OsiriX MD, version 12.0.3). New lesions identified after SIV infection were called “relapsed lesions” and assumed to be disseminated from pre-existing sites of disease. CD4 counts and SIV viral RNA copies from plasma and tissues were also measured as previously published (17).

To determine whether the severity and distribution of disease involvement prior to treatment were associated with relapse, spider plots of each animal were generated by connecting the locus of points created by plotting a set of PET CT-based disease measurements on a polar axis (Python 2.7.15, Matplotlib 2.2.3). The angular coordinates correspond to the metric and the radial coordinate gives the value of the metric normalized to the maximum value of that metric in the dataset. The maximum value of each metric was determined across all animals and timepoints. The value of the radial coordinate is the value of the metric measured in that specific animal at the specific timepoint divided by the maximum value of that metric across all animals and timepoints.

The measurements included: total lung FDG activity (square-root-transformed, 0 – 695.312908), number of extrapulmonary sites (0 – 5), lung lobes involved (1–7), necrotic thoracic lymph nodes (0 – 6), FDG-active thoracic lymph nodes (0 – 13), lung cavity (0=no, 1=yes), granuloma clusters (0=no, 1=yes), consolidation (0=no, 1=yes), bilateral lung disease (0=unilateral, 1=bilateral).

### Flow Cytometry Analysis

Flow cytometry was performed on PBMC, BAL and tissues using a combination of antibodies (Supplementary Table 2A), as described(17). PBMC and BAL cells were stimulated with Mtb ESAT6-CFP10 overlapping peptide pools (10µg/ml of each peptide) and/or only media (RPMI+10%hAB) prior to staining. Data acquisition using an LSR II (BD) was performed and FlowJo Software v.9.7 (Treestar Inc, Ashland, OR) was used for analysis. The gating strategy for cytokines and cytotoxic markers is shown in Supplementary Figure 1.

### Circos Plots

For each animal, the identified bar-codes in each tissue and associated metadata including tissue anatomical location and time of lesion detection (pre-and post-SIV infection) were converted into text formats using custom scripts compatible with plotting by Circos software(20). Plots were manually adjusted such that links between tissues sharing bar-codes were drawn, when possible, to originate from lung sites detected early by PET CT with the highest CFU burden; however, other lung sites with the same bar-code may also be the source of dissemination between tissues. Custom scripts and relevant configuration files used to generate the Circos plots are available upon request.

### Statistical analysis

Shapiro-Wilk test was used to test for normality. Mann-Whitney test was used for non-paired data and Wilcoxon matched-pairs signed rank test was used for any paired data for comparing two groups. For longitudinal PBMC and BAL data, groups were compared at each time point with Mann-Whitney tests (not adjusted for multiple comparisons). Longitudinal PET data was analyzed using a mixed-effects model fit using Restricted Maximum Likelihood (REML) with Dunnett’s multiple comparisons test adjusted p-values reported. Categorical data was analyzed using Fisher’s exact test. For counts (including cell counts, CFU, and FDG activity), the data was first transformed (adding 1 to entire dataset), so that zeroes could be visualized and analyzed on a log_10_ scale. All statistical tests are two-sided and significance was established at p £ 0.05. Figures, graphs and statistical tests were performed in GraphPad Prism Mac OSX (Version 10.1.1, GraphPad San Diego, CA) or JMP® Pro 17.2.0.

### Ethics statement

All animal procedures were approved by the University of Pittsburgh Institutional Animal Care and Use Committee (protocol 14023305) in compliance with the Animal Welfare Act and Guide for the Care and Use of the Laboratory Animals.

### Funders

This work was supported by the Otis Childs Trust (P.L.L.) and National Institutes of Health (NIH R01AI111871[P.L.L.] and AI134195[P.L.L.]) who had no role in the design, data analysis/interpretation, or manuscript construction.

## Results

### PET CT defined relapse is associated with higher bacterial burden

Macaques with clinically defined active TB disease have a range of radiographic patterns that are seen in human TB including: cavities (e.g., 16814, Figure 1B, green arrow), granulomas of various size, multiple lung lobe involvement, lung consolidation (e.g., 16414 pre-drug image, Figure 1B, purple arrow), tree-n-bud pattern, lymphadenopathy with or without necrosis, etc. The standard treatment of drug-susceptible TB includes 2 months of rifampin (RIF), isoniazid (INH), pyrazinamide, and ethambutol, followed by 4 more months of RIF and INH. As this was pilot study for model development with a limited number of animals (n=12), a short course of treatment with only RIF and INH (the two backbones of standard TB treatment) for 2 months was chosen to prevent complete sterilization of Mtb and increase the likelihood of relapse. Following treatment, macaques had a 1 month, drug-free period and then SIV infection (Figure 1A). During drug treatment, there was an overall reduction in lung inflammation (measured as total lung FDG activity), but no difference in change of inflammation over time between animals that would later develop relapse compared to non-relapse animals (Supplementary Figure 2A). While inflammation of individual lymph nodes varied in animals over time, on average, thoracic lymph node inflammation (measured as standard uptake volume ratio, SUVR) decreased during treatment (fold change mean: no-relapse 0.95, relapse 0.61) (Supplementary Figure 2B). Lesion-specific reductions in size and inflammation were also observed, especially in large consolidations and cavities (Figure 1B). Only one animal (31216, relapsed) had a persistent consolidation but with reduced size and metabolic activity during drug treatment while the remaining animals had near resolution of consolidations on scan. Lesional changes associated treatment varied over time (i.e., some lesions took longer to improve than others) and were independent of bacterial burden at necropsy (Supplementary Figure 2C) as expected given the heterogeneity in bacterial burden and host response in individual granulomas(21–23).

Relapse TB was defined as the development of a new lung granuloma detected by PET CT (and confirmed at necropsy) after SIV infection (Figure 1B, Supplementary Figure 3A). Eight of the 12 animals developed relapse. Four of the eight relapse animals had microbiologic and/or clinical evidence of disease (i.e., tachypnea and Mtb detected); non-relapse animals had no such signs (Figure 2). At necropsy, relapse macaques had greater pathology and more Mtb CFU (95% CI of log_10_ difference between medians 3.8 to 6.3) than non-relapse animals, particularly in the lung and thoracic lymph nodes (Figure 3, Supplementary Figure 3B-D). Serial PET CT images allowed us to track granulomas that appeared before and after treatment. Relapse animals had a significantly higher frequency of granulomas (median of 68%) with viable Mtb at necropsy. Among the new granulomas that appeared during relapse, 84% had viable Mtb growth. In contrast, only one of the four non-relapsed animals had Mtb growth at necropsy in a single granuloma (184 CFU) (Supplementary Figure 3E-F). Thus, while killing all viable Mtb in every anatomic reservoir would be an optimal strategy for drug treatment, some viable Mtb are likely to persist after “successful” treatment emphasizing the importance of on-going immune control in preventing relapse.

**Figure 2.**
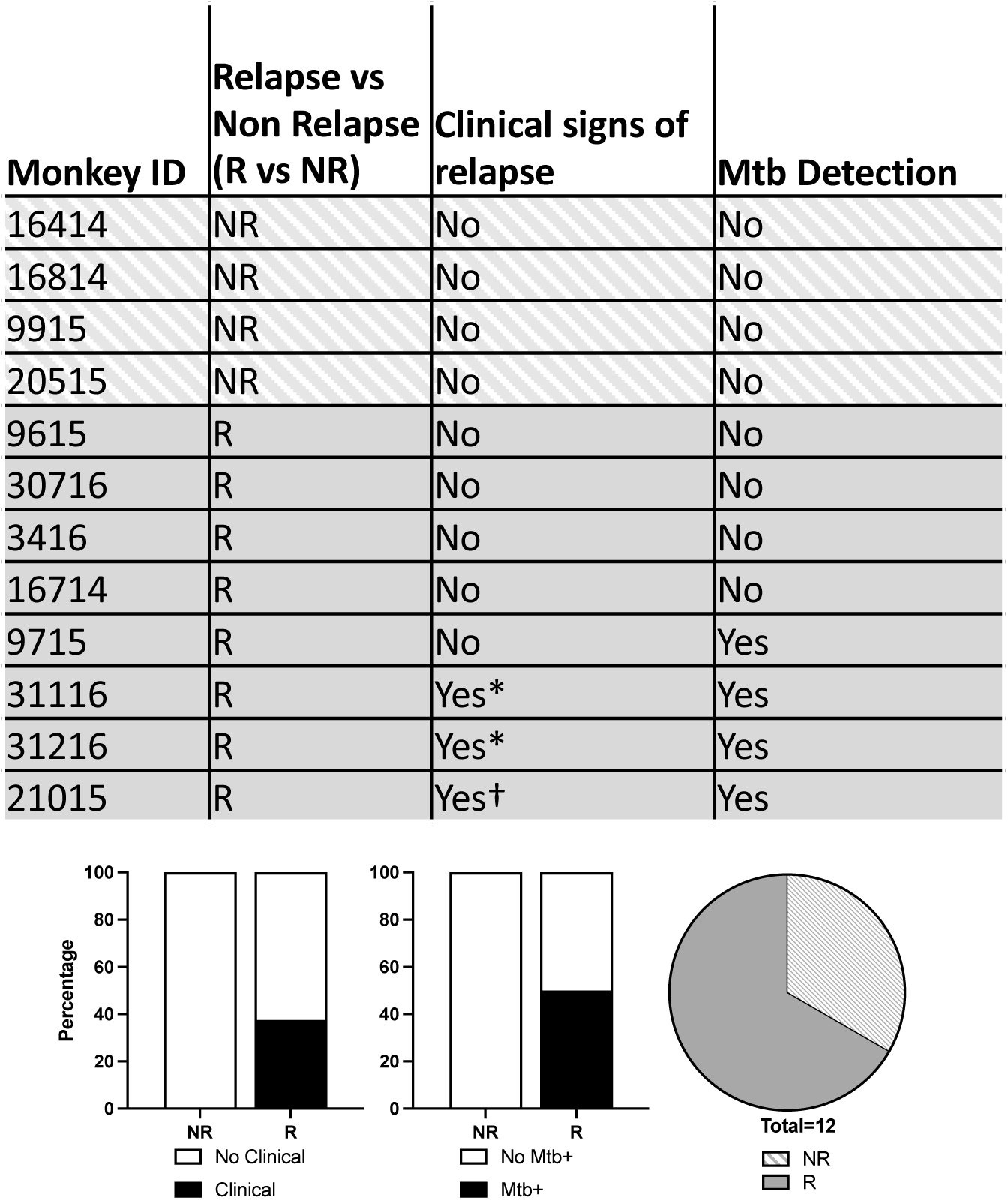
Clinical and microbiological evidence of relapse after SIV infection among radiologically defined relapse and non-relapse animals. Mtb detection refers to microbiologic evidence (Mtb detected) in either gastric aspirate (GA) or bronchoalveolar lavage (BAL) samples *Increased respiratory effort. Irregular respirations. Bar graphs represent percentage of animals in each outcome group with clinical or microbiologic evidence of relapse post-SIV infection.

**Figure 3.**
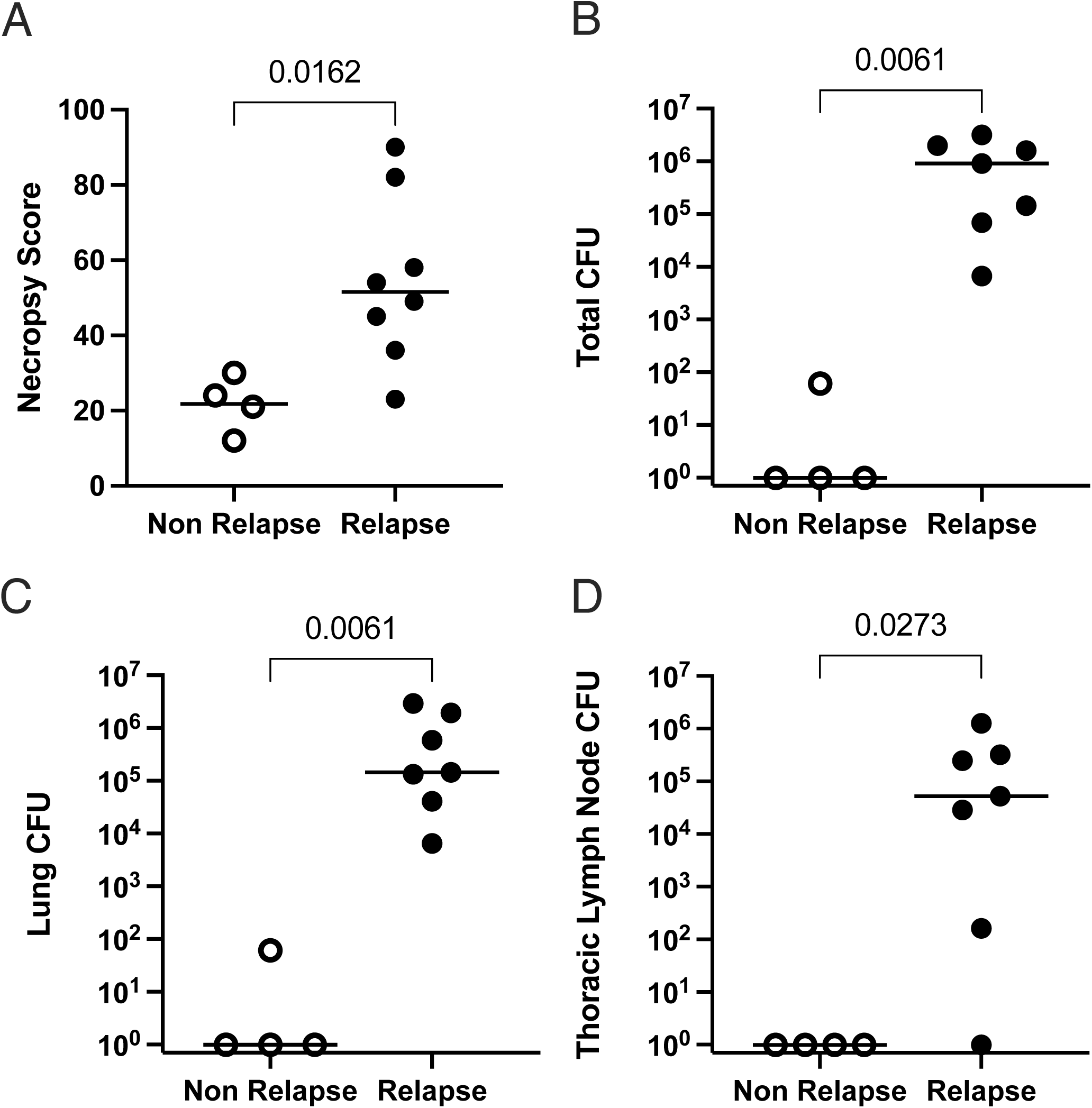
PET CT defined relapse is associated with greater gross pathology at necropsy (measured by necropsy score) (A), total Mtb burden (measured as CFU, colony forming units of Mtb) (B), lung Mtb burden (C) and thoracic lymph node burden (D) compared to non-relapse animals. P-values shown were determined by Mann-Whitney test. Each dot represents an animal, and lines shown are medians.

### Pre-treatment radiographic characteristics are not associated with relapse risk

We sought to determine whether PET CT characteristics before TB treatment were associated with relapse. We previously published that total lung inflammation (total lung FDG activity) correlates with bacterial burden(19) and this measure was similar between relapse and non-relapse animals before treatment (Figure 4A). To determine whether the extent of lung or lymph node involvement prior to treatment was associated with relapse, we compared PET CT features of disease severity including: number of lung lobes involved, presence of bilateral lung involvement, consolidation (defined as a lesion greater than 2 cm), clusters of TB lesions, cavity, metabolically active thoracic lymph nodes, and extrapulmonary disease (Figure 4B-H). All these parameters were similar between relapse and non-relapse groups as were the reductions in the total lung FDG and lymph node FDG (Supplementary Figure 2). To examine the combined effect of disease severity, spider plots of each feature were generated with a score based on the extent of each disease feature. Scores were similar between groups before SIV induced relapse (Supplementary Figure 4). Not surprisingly, the severity score was higher in relapse animals at necropsy (Supplementary Figure 4D).

**Figure 4.**
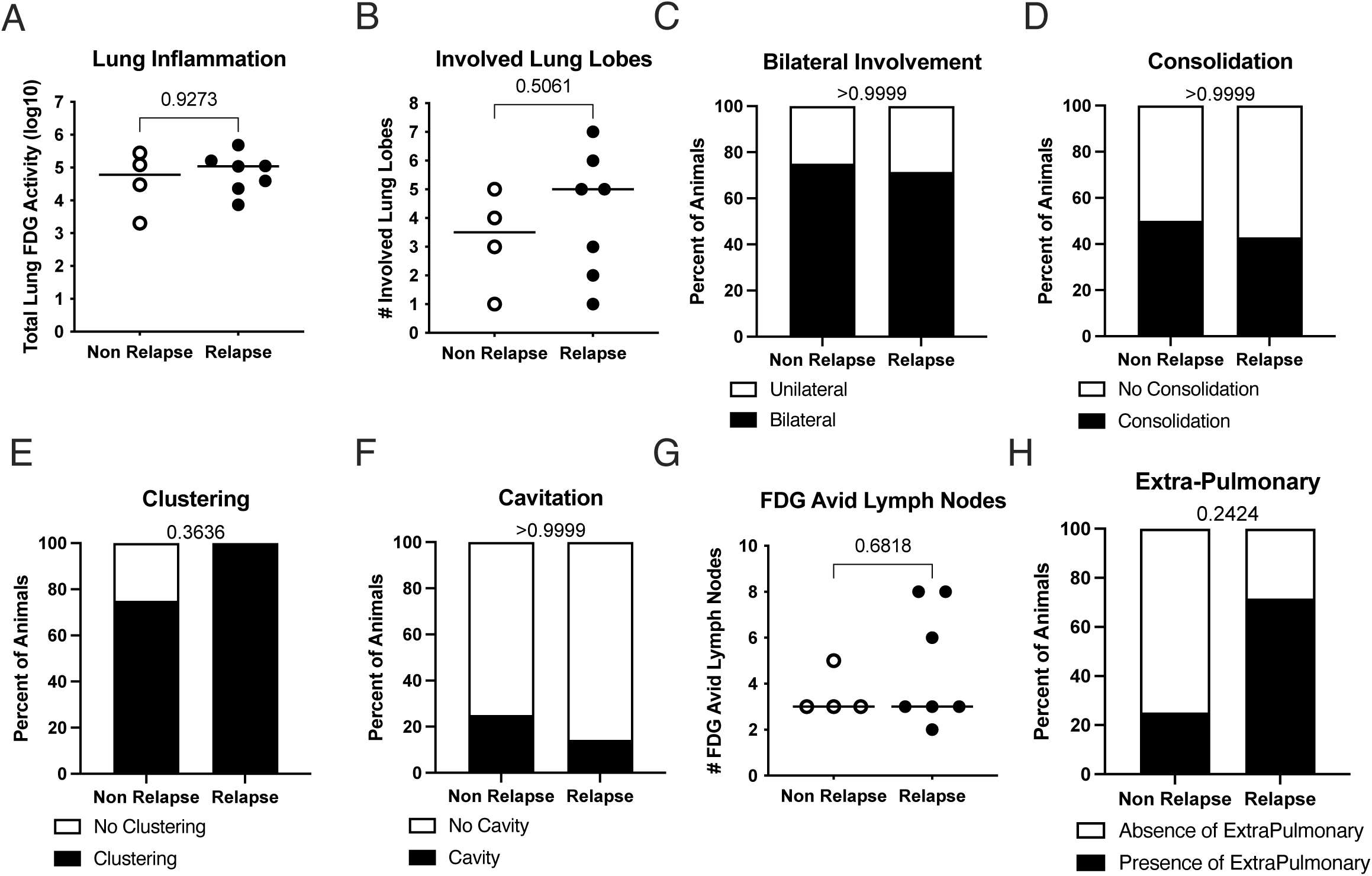
PET CT characteristics prior to TB drug treatment. PET CT characteristics were similar between non-relapse (n=4) and relapse (n=7; one animal could not be scanned prior to drug-treatment) animals that included: total lung inflammation (measured using FDG activity) (A), number of involved lung lobes (B), proportion of unilateral or bilateral lung involvement (C), proportion of animals with lung consolidation as a CT feature (D), proportion of the animals with clusters of granulomas as a feature of TB disease on PET CT (E), proportion of animals with cavitary disease on PET CT scan (F), number of FDG-avid (as a marker of inflammation) mediastinal lymph nodes (G), and proportion of animals with PET CT findings of extrapulmonary disease prior to drug treatment (H). P values are based on Mann-Whitney (A, B, G) or Fisher9s Exact (C, D, E, F, H) tests.

Given that our prior data showed that TB granulomas are heterogeneous within the same host and can independently influence disease risk(21–24), we examined individual granuloma characteristics. Maximum granuloma size and/or maximum SUV (metabolic avidity) per animal before treatment and/or after treatment was not associated with relapse (Supplementary Figure 5). We then took advantage of serial PET CT scanning data in these animals to examine whether other unique PET CT features such as resolution of granulomas were associated with relapse risk. Interestingly, all 4 of the non-relapse animals had at least 1 lung granuloma resolve completely (defined as the absence of lesion seen by PET CT on at least two subsequent scans and not found at necropsy) on PET CT but only one of eight relapse animals had lesion resolution (100% vs 12.5%, p<0.01, Fisher’s Exact).

### Distribution of bar-codes during relapse

We previously established that each individual granuloma is established by a single bacillus, distinguishable by a molecular bar-code, whereas thoracic lymph nodes often have multiple bar-codes due to migration of Mtb from lung granulomas(16, 22, 25). Accordingly, bar-coded Mtb libraries allow us to track dissemination of individual Mtb bacilli. At necropsy, we harvested PET CT scan-matched lesions from lung, thoracic lymph nodes and other tissue sites while discriminating whether they appeared before SIV infection (or “pre-SIV”) or during relapse (or “post SIV”). Bar-codes recovered from scan-matched granulomas appearing after SIV infection were mapped to both thoracic lymph nodes and lung granulomas identified before SIV infection (Figure 5A, Supplementary Figure 6,7). We found that relapse could be dominated by a single bar-code strain (monkey 3416) or multiple strains. In the case of animal 31216, one of the bar-codes identified from a relapsed granuloma could be matched only to a pre-treatment thoracic lymph node. These data emphasize the importance of eradicating all viable Mtb from both lung and lymph nodes. Eliminating viable Mtb within lymph nodes is particularly important because they exhibit reduced bacterial killing during drug treatment(26). However, not all bar-codes recovered from pre-SIV lesions (i.e., existed during drug treatment) disseminated to relapse sites, indicating that not all viable Mtb lesions after drug treatment were capable of dissemination even during SIV infection. In fact, only 42% (median, range 0-100%) of bar-codes from pre-treatment lung granulomas were observed in relapse granulomas, highlighting the independent risk of each granuloma to disseminate. Lastly, we investigated the proximity of new granulomas to the (“pre-SIV”) originating granuloma or lymph node using bar-code data and XYZ coordinates from PET CT scans before and after drug treatment (Supplementary Figure 6,7). New granulomas during relapse appeared to be closer in distance to a thoracic lymph nodes than to old granulomas matched by bar-code (Figure 5B).

**Figure 5.**
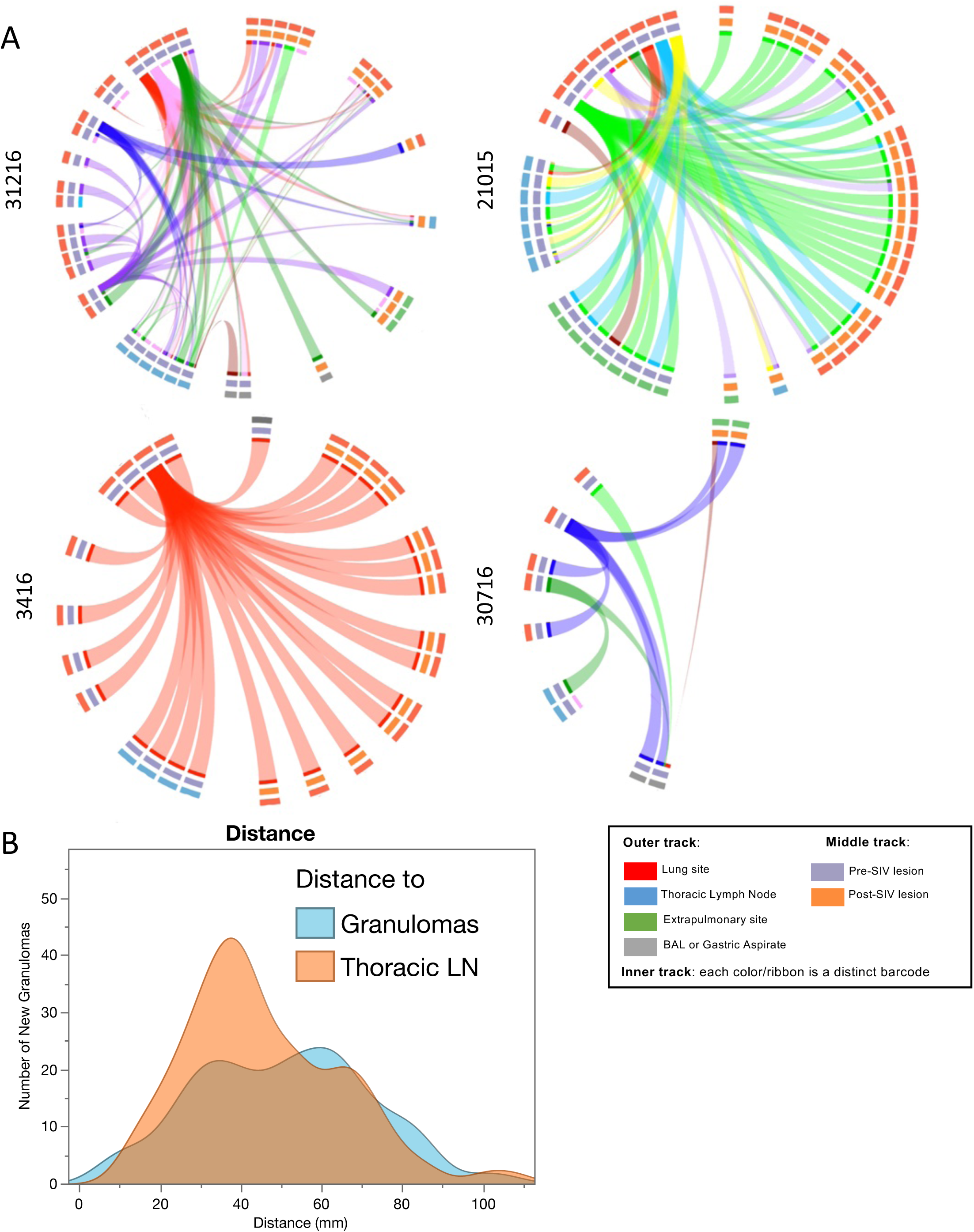
Distribution of barcodes by anatomical compartment. (A) Circos plots representing barcoded Mtb detected in tissues (inner track), with ribbons representing tissues containing the same barcodes. Middle track represents timing of sample seen on scans (pre-or post-SIV infection) and outer track represents the anatomical compartment associated with each tissue. Distinct barcodes are represented by a different color. (B) New granulomas appear to be closer in distance to founding (seen at 4 weeks post-Mtb infection) lymph nodes (green) than founding granulomas (blue). Y-axis shows the number of granulomas sharing barcodes with founding granulomas and lymph nodes and x-axis shows distance between the new granuloma and founding granuloma/lymph node.

### Higher SIV RNA levels observed in relapse granulomas and lymph nodes though immune responses did not distinguish relapse outcomes

We found that plasma SIV viral loads were similar between relapse and non-relapse animals during the course of SIV infection as were peripheral blood CD8 and CD4 T cells (Supplementary Figure 8A,B). In general, neither lung nor systemic immune responses distinguished relapse and non-relapse cases. Granuloma-specific Th1 cytokine responses among CD4 and CD8 T cells were similar between relapse and non-relapse animals (Supplementary Figure 9). Th1 (IFN-γ, IL-2, TNF), IL-10, and IL-17 T cell responses were also similar in the airways though animals that would later develop relapse had lower IL-10 production among alveolar macrophages before SIV infection (Supplementary Figures 10-11). In the blood, higher frequencies of terminally differentiated (CD27-/CD45R+) CD4 and CD8 T cells were observed in the non-relapse animals at very early timepoints after Mtb infection (Supplementary Figure 12) but not later. Blood immune responses were similar between relapse and non-relapse animals (Supplementary Figure 13-20). Higher SIV RNA levels were observed in granulomas with viable Mtb compared to those without detectable Mtb and in the non-granulomatous lymph nodes (avoiding cellular distortion from granulomas) of relapsed animals compared to non-relapse animals (Supplementary Figure 8C). These data are consistent with our prior work showing discordance between systemic and localized immune responses(23) as well as the synergistic relationship between SIV and Mtb within tissues(17).

## Discussion

Using our macaque model, we developed a model of TB relapse and characterized the events during the treatment of active TB and SIV-induced relapse. While relapse was defined as a newly developed lesion after drug treatment, it was validated by gross pathology and bacterial burden at necropsy. We found that PET CT characteristics before drug treatment (Figure 4) did not correlate with relapse outcome. And changes in inflammation (FDG avidity) or size of individual lesions during treatment were not associated with the presence or absence of viable Mtb at necropsy nor with the likelihood of dissemination occurring from a specific lesion. Our findings are consistent with human TB treatment trials in which no PET CT characteristics before, during, or at the end of shortened treatment could definitively predict relapse and variable responses in size and metabolic activity of different lesion types were observed in the same host (reviewed in (27)). To our knowledge, this is the first study to track bacterial dissemination during SIV-induced TB relapse. Using bar-coded Mtb, we were able to harvest scan-matched lesions, correlate the timing of appearance during infection, treatment and post-SIV, and track dissemination. Only 42% of bar-codes from “pre-SIV” lesions could be detected in relapse lesions demonstrating that not all granulomas with viable Mtb are capable of dissemination, even during acute SIV infection. This is consistent with our findings that granulomas function independently from one another in the same host(23, 24). Bar-codes found in the relapse lesions could be traced to both lung granulomas and thoracic lymph node sites observed prior to SIV infection. In one case, a bar-code from a relapsed lesion could only be traced to a thoracic lymph node underscoring the importance of lymph nodes as a reservoir for relapse. Lastly, one animal without relapse still had a low burden of Mtb in a granuloma. Thus, it stands to reason that the primary goal of TB treatment should be rapid and complete sterilization from all anatomic reservoirs even though some low-level persistence of Mtb may not result in clinical relapse. This is consistent with human studies where sputum or airway Mtb RNA occurred in the context of “non-resolving” or “intensifying” PET CT lesions observed at the conclusion of standard TB treatment, without evidence of relapse noted at the one-year follow-up period(28).

To date, there are no reliable biomarkers that accurately predict TB cure making efforts to shorten TB treatment a significant challenge. For example, in a recent 4-month treatment trial, relapse rates were unacceptably high (∼20%) forcing the study to be stopped prematurely(29). Biomarkers associated with TB cure including systemic immune markers (e.g., CD27, plasma cytokines) and microbiologic markers (e.g., sputum conversion) have had limited success (reviewed in (5)). PET CT studies vary in patient population, drug regimen, immune status (HIV-infected vs HIV-naïve), interval assessments after treatment and imaging metrics(27). Using PET CT to monitor the treatment of drug susceptible TB patients, 14% of patients had a resolved pattern (no metabolic activity) but the remaining patients had either persistent metabolic activity or a mixed pattern (increased metabolic activity or new lesions). While no failures were observed in the resolved group, but treatment failure was observed in 28% of those with a mixed pattern(10). The risk of treatment failure was higher among patients who had less than an 80% reduction in total glycolytic activity index (a measure of lung metabolic activity via FDG) from the time of diagnosis to month 6 of treatment(10). Other studies focused on patients with HIV-TB coinfection have shown that using the maximum metabolic activity of involved lymph nodes (both pulmonary and extrapulmonary) was a better predictor of microbiologic response to therapy(27). Overall, it seems that the absence of residual lung metabolic activity by PET at the end of treatment (independent of HIV status) appears to be associated with cure but residual metabolic activity is common and without predictive value for relapse (reviewed in (27)).

There are several limitations to this study. We purposely chose a short course of RIF and INH to increase the probability of relapse as proof of principle given the limited number of animals in this study. Since we had not performed relapse studies previously in this model, we were unable to perform a robust power analysis to determine an adequate sample size. Given the human relapse data, it is probable that large numbers of animals would be needed, which is neither practically nor ethically feasible. Nonetheless, the short course treatment was effective in some animals providing the opportunity to compare relapse and non-relapse in this study. Importantly, this study does not replicate the more common scenario of HIV-TB coinfection in which Mtb burden is likely to be much higher prior to TB treatment and where immune driven efforts to sterilize Mtb are hindered by HIV-driven immune suppression.

In summary, we have developed a model of TB relapse in which the source of bacterial dissemination can be tracked and PET CT characteristics can be validated by bacterial burden at necropsy. This model could be useful in assessing new short course treatment regimens and for gaining insights into the events that occur during relapse.

## Acknowledgements

Special thanks to Elise Chu for the assistance in the figure design. We thank the Ambrose, Lin, Flynn and Fortune labs for their veterinary and technical dedication as well as the University of Pittsburgh’s Division of Laboratory Animal Resources.

---

**Supplementary Figure 1.**
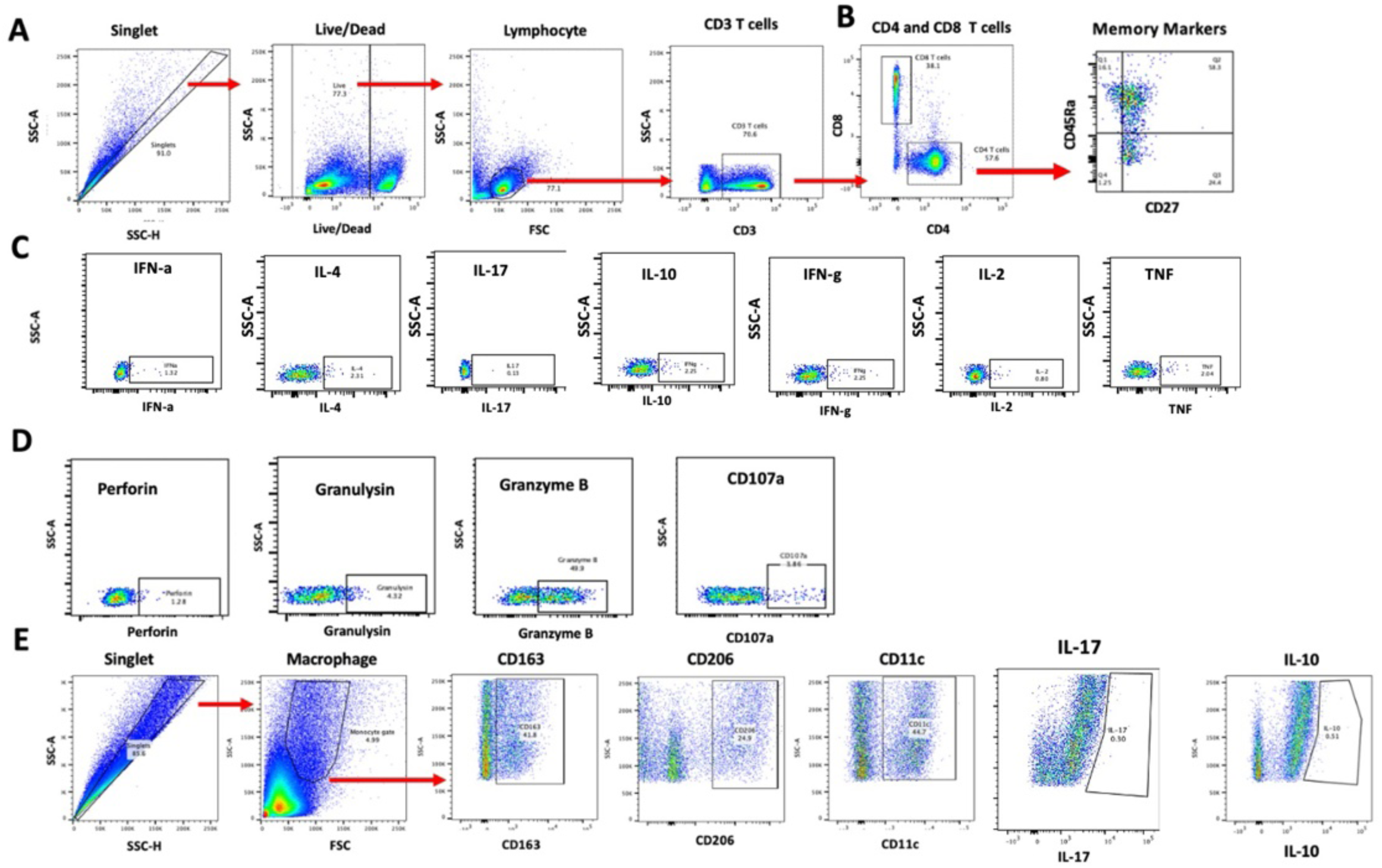
Example gating strategy for flow cytometry. T cell gating strategy is the same for PBMC, BAL, and tissue samples. Macrophage gating is specific to BAL cells. A) PBMC stimulated with PDBU and Ionomycin. Singlets were positively selected. Live cells were selected out of the single cell gate. Lymphocytes were selected from the live cell gate. CD3 T cells were selected from lymphocytes and CD4 and CD8 T cells were selected from CD3 T cell gate. B) Memory Markers were selected from either CD4 or CD8 gate in PBMC. C) IFN-a, IL-4, IL-17, IL-10, IFN-g, IL-2, TNF were identified from the CD3 T cell gate. D) Perforin, Granulysin, Granzyme B, and CD107a were identified in CD3 T cells. E) Singlets were positively selected from BAL. Macrophage gate was selected from the singlet gate. CD163, CD206, CD11c, IL-17, and IL-10 from the monocyte gate.

**Supplementary Figure 2.**
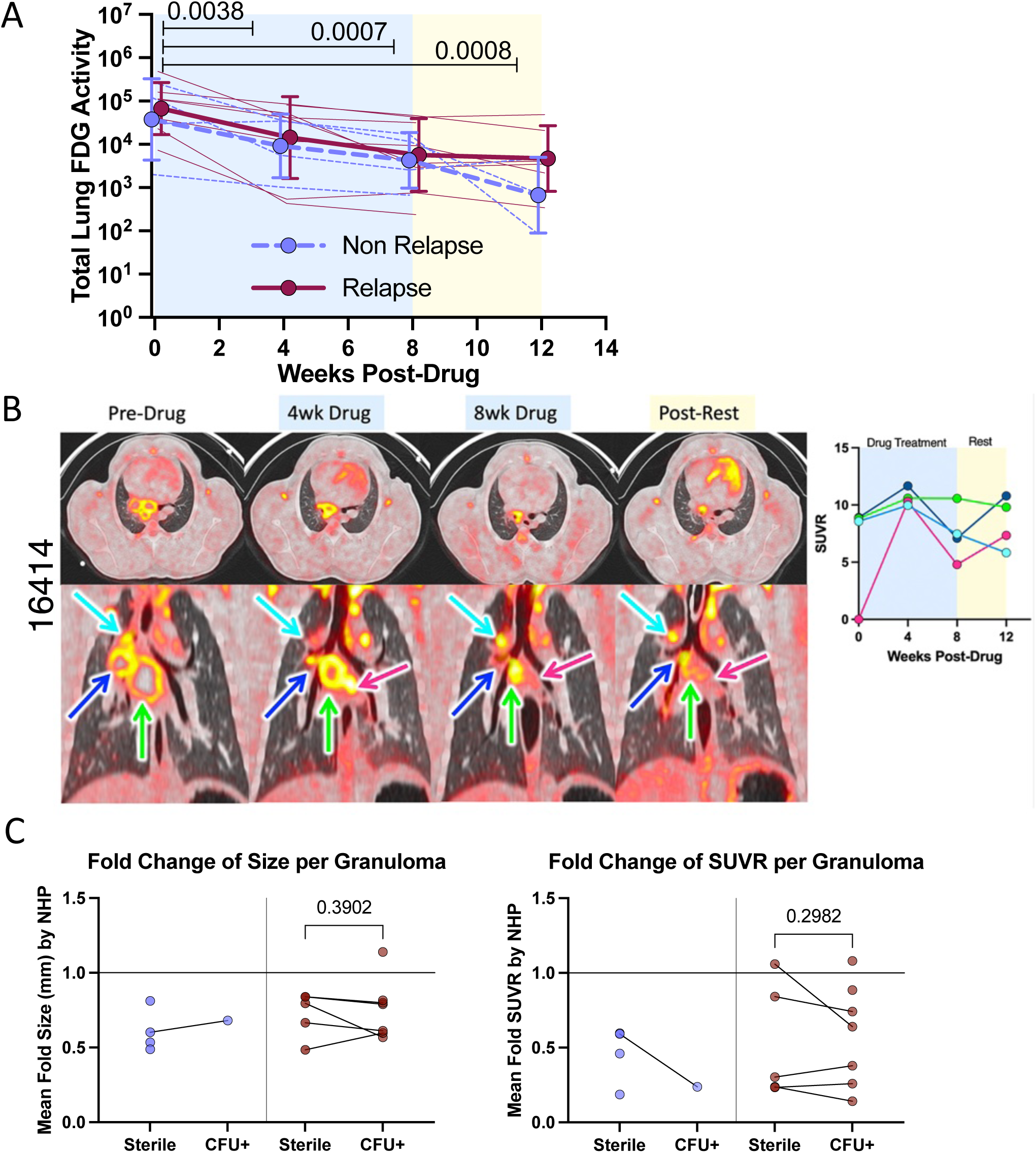
(A) Lung inflammation (FDG). A mixed-effects model showed that there was no difference between the two groups; however, FDG significantly decreased after drug treatment. Darker lines show mean and SD for each group; lighter lines represent trajectory of each animal. P values shown represent Dunnett9s multiple comparison adjusted values comparing each time point to pre-drug treatment. **(B)** Independent and dynamic changes in metabolic activity among TB-associated thoracic lymph nodes (LN) before, during and after TB drug treatment. Axial (top row) and coronal images (lower row) of thoracic LN are shown. All lymph nodes increase in metabolic activity (standard uptake value ratio, SUVR) between 0 and 4 weeks-treatment, but diverge in pattern over time. Color-matched thoracic LN are shown in the coronal views (bottom row) and line plots (right) over time. **(C)** Change in granuloma size and avidity during drug treatment does not reflect bacterial burden. (Left) Average fold changes in size (mm) in sterile and CFU+ granulomas matched per animal. (Right) Average fold changes in avidity (SUVR) in sterile and CFU+ granulomas matched per animal. Each dot is the mean of all changes of granulomas per animal of either sterile (left) or CFU+ (right) granulomas. Lines connect averages per each individual animal. Only one animal in the non-relapse group had a granuloma that grew CFU. Two animals in the relapse group had no sterile granuloma that could be confidently analyzed on PET CT.

**Supplementary Figure 3.**
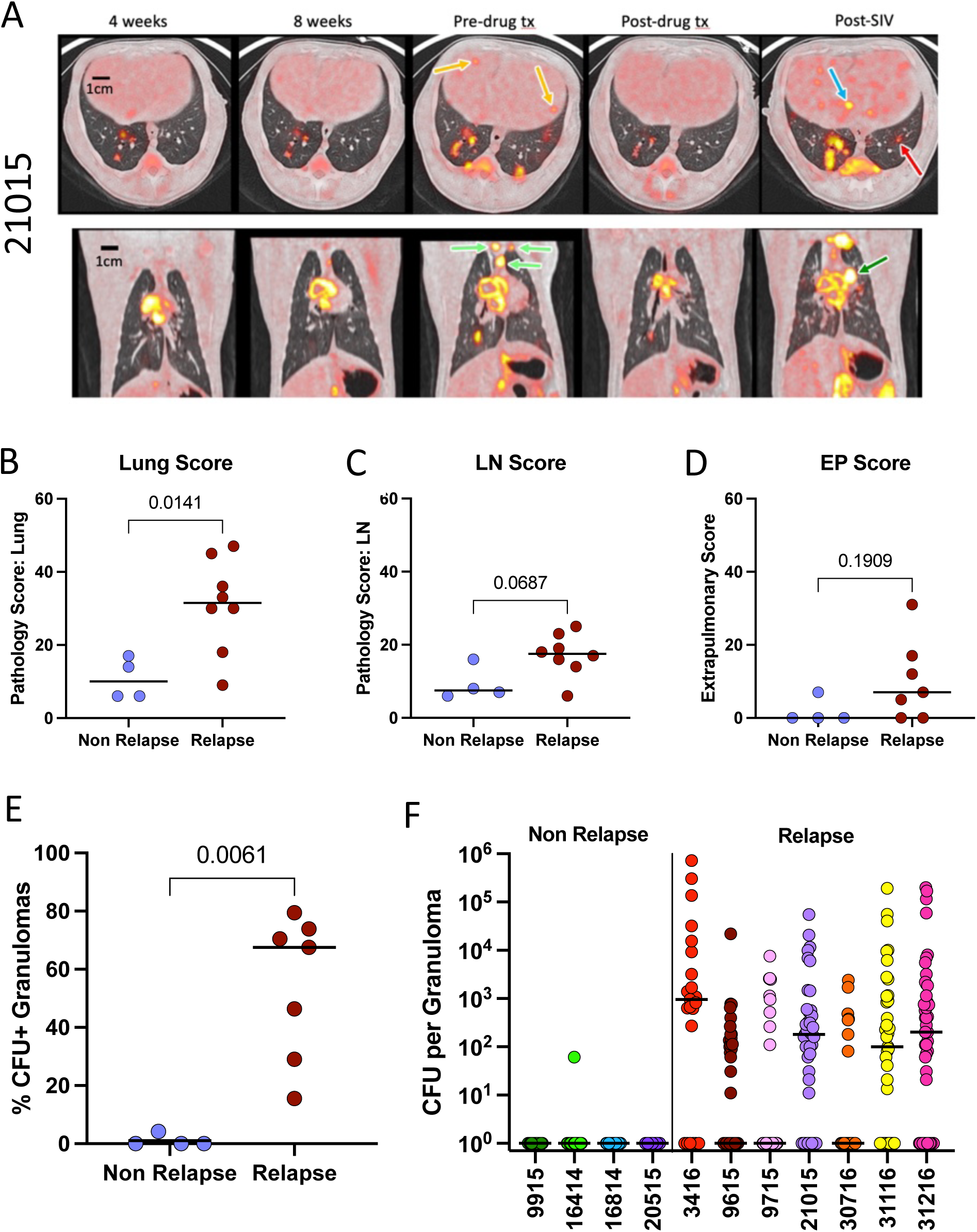
(A) PET CT evidence of SIV-induced relapse after TB drug treatment. Top row: Synchronized axial images show liver granulomas before drug treatment (orange arrows) that resolve on subsequent scans, but new granulomas are seen post-SIV (blue arrow). While granulomas and clusters can be seen on all scans prior to SIV, there is a new granuloma (red arrow) in the left lower lobe. Bottom row: Coronal images show thoracic lymph node involvement in the upper mediastinum before drug treatment (light green arrows) and new post-SIV (dark green arrow). TB involvement of post-SIV lymph node was confirmed at necropsy. (B) PET CT defined relapse is associated with higher lung pathology and a trend to higher thoracic lymph node (LN) pathology (C). (D) Extrapulmonary score (derived from gross pathology and Mtb growth) was similar between relapse and non-relapse animals. (E) Relapse animals had a greater proportion of lung granulomas with Mtb growth. (F) Relapse animals have greater bacterial burden per lung granuloma (measured as colony forming units, CFU) compared to non-relapse animals. P-values shown reflect Mann-Whitney test. Each dot represents an animal, and lines represent medians.

**Supplementary Figure 4.**
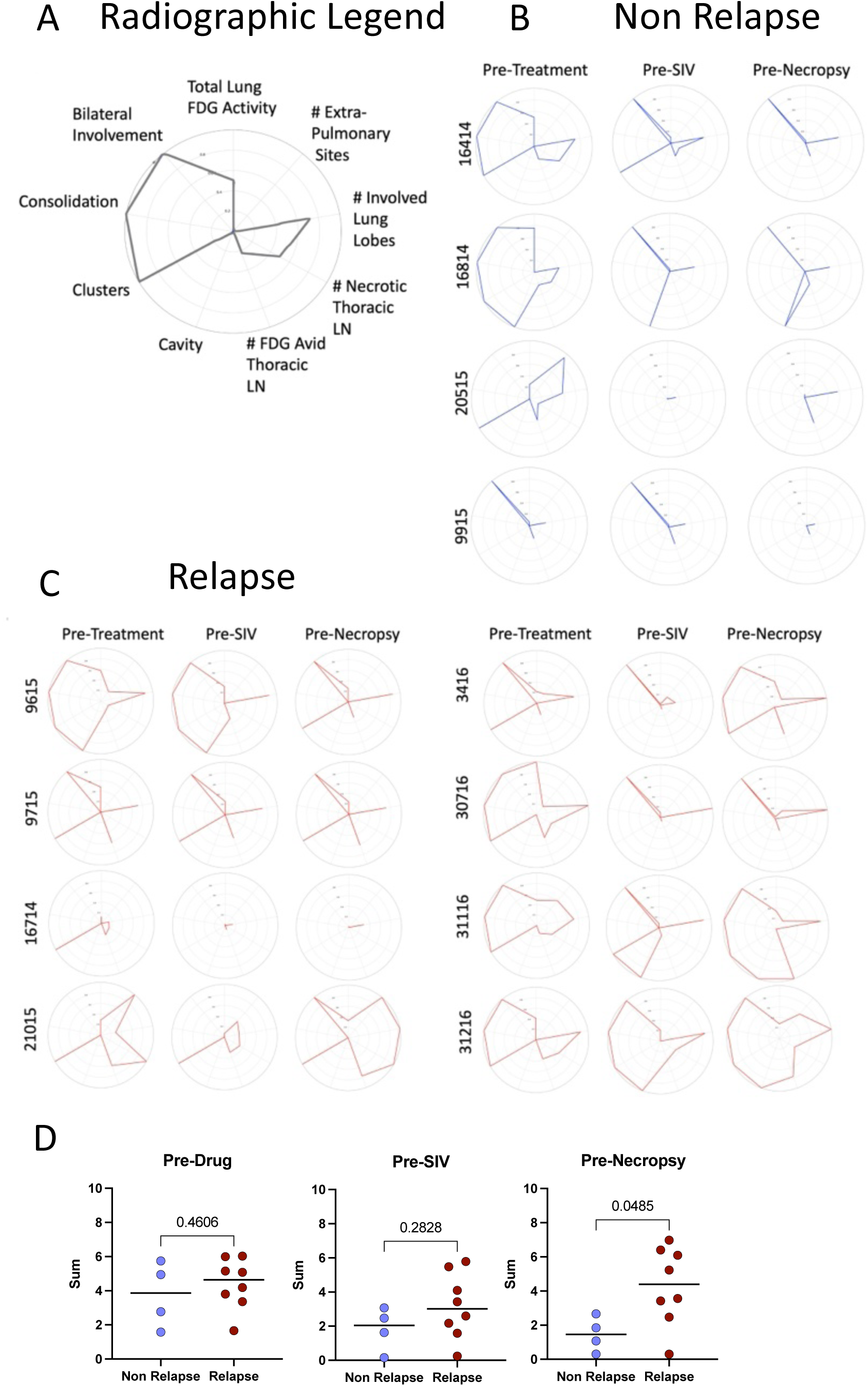
(A) Spider plot shows the radiographic features of TB disease in composite with each radial coordinate measuring the severity of involvement. Composite of PET CT identified features of tuberculosis prior to treatment, after TB drug treatment but before SIV infection, before necropsy among animals that (B) did not relapse (B) and those that did relapse (C). FDG: F^18^ Fluoro-DeoxyGlucose, EP: Extra-Pulmonary, LN: Lymph Nodes. D) The sum of the radiographic feature scores were compared between relapse and non-relapse animals at serial time points Each dot represents an animal, lines represent medians. Mann-Whitney p-values shown.

**Supplementary Figure 5.**
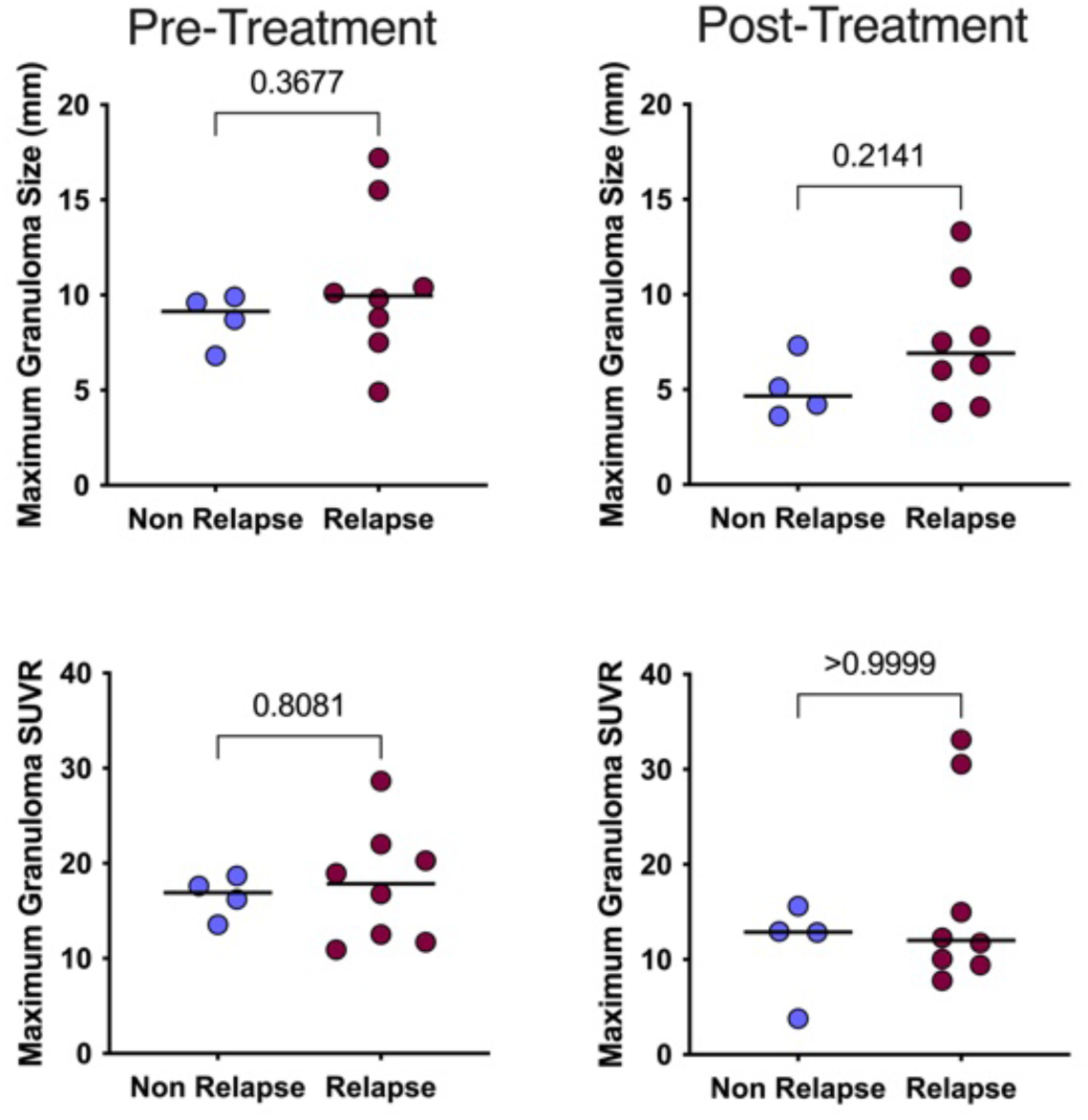
PET CT features before and during drug treatment among relapsed (N=8) and non-relapsed (n=4) animals. Maximum size and metabolic activity (measured standard uptake value ratio, SUVR) for each animal before and after drug treatment. Each dot represents the maximum value for each animal, lines represent medians. P-values reflect Mann-Whitney analysis.

**Supplementary Figure 6.**
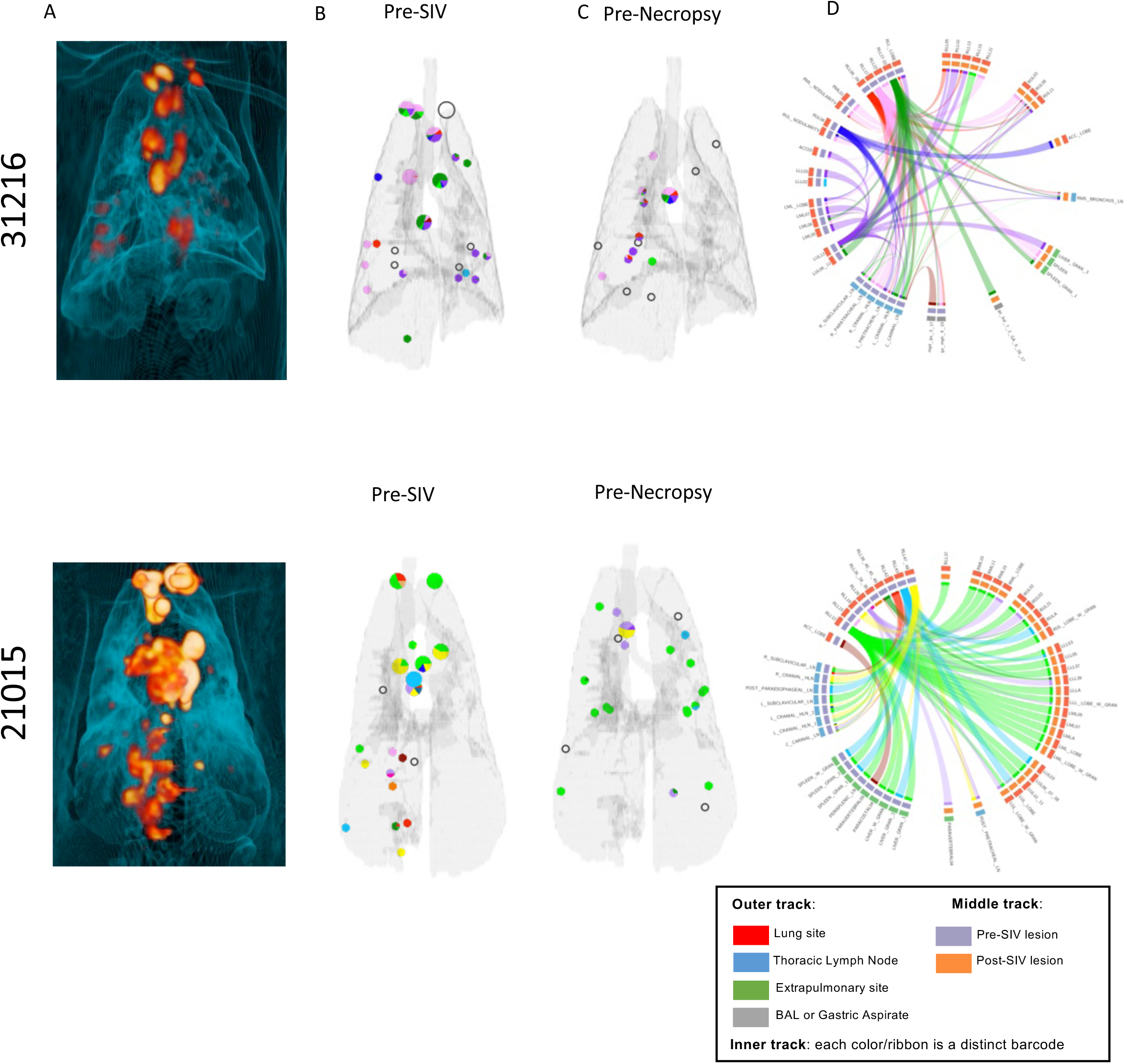
Barcode distribution of disease for animals 31216 (top row) and 21015 (bottom panel). A) Three-dimensional volume rendering of lung disease pre-necropsy scan. (B) Anatomic distribution of Mtb barcodes from pre-existing granulomas and thoracic lymph nodes seen on scan before SIV-infection (B) and new granulomas and lymph nodes observed after SIV infection (pre-necropsy) (C). Open circles represent sterile tissue (B and C). Smaller circles represent granulomas or clusters and larger circles represent thoracic lymph nodes. (D) Circos plots representing barcoded Mtb detected in tissues (inner track), with ribbons representing tissues containing the same barcodes. Middle track represents timing of sample seen on scans (pre-or post-SIV infection) and outer track represents the anatomical compartment associated with each tissue. Distinct barcodes are represented by a different color, matched across B, C, and D.

**Supplementary Figure 7.**
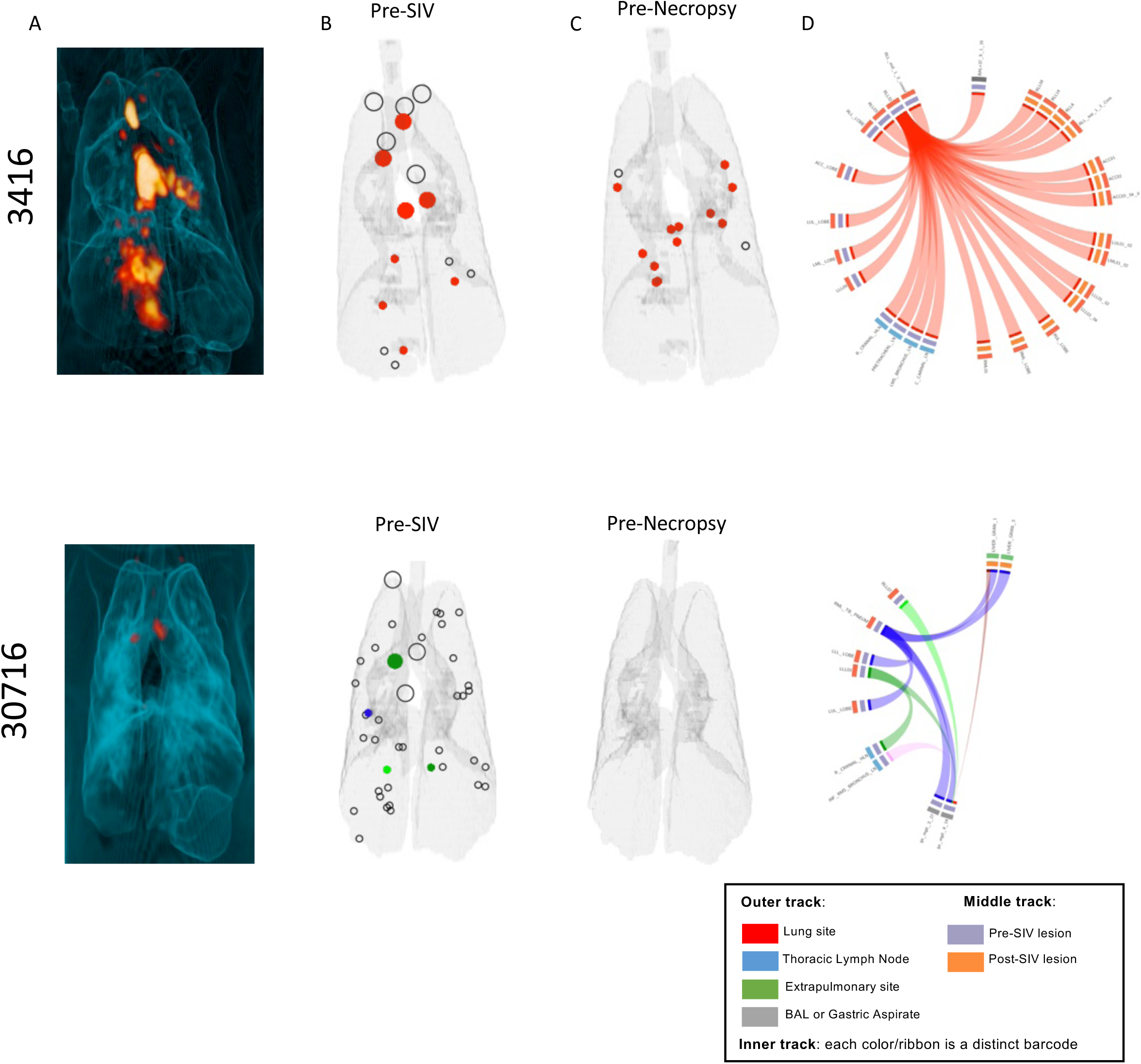
Barcode distribution of disease for animals 3416 (top row) and 30716 (bottom panel). A) Three-dimensional volume rendering of lung and lymph node disease on pre-necropsy scan. (B) Anatomic distribution of barcodes found in granulomas and thoracic lymph nodes seen on scan before SIV-infection. (C) Distribution of new granulomas and lymph nodes seen on scan after SIV-infection (pre-necropsy). Open circles represent sterile tissue (B and C). Smaller circles represent granulomas or clusters and larger circles represent thoracic lymph nodes. (D) Circos plots representing barcoded Mtb detected in tissues (inner track), with ribbons representing tissues containing the same barcodes. Middle track represents timing of sample seen on scans (pre-or post-SIV infection) and outer track represents the anatomical compartment associated with each tissue. Distinct barcodes are represented by a different color, matched across B, C, and D.

**Supplementary Figure 8.**
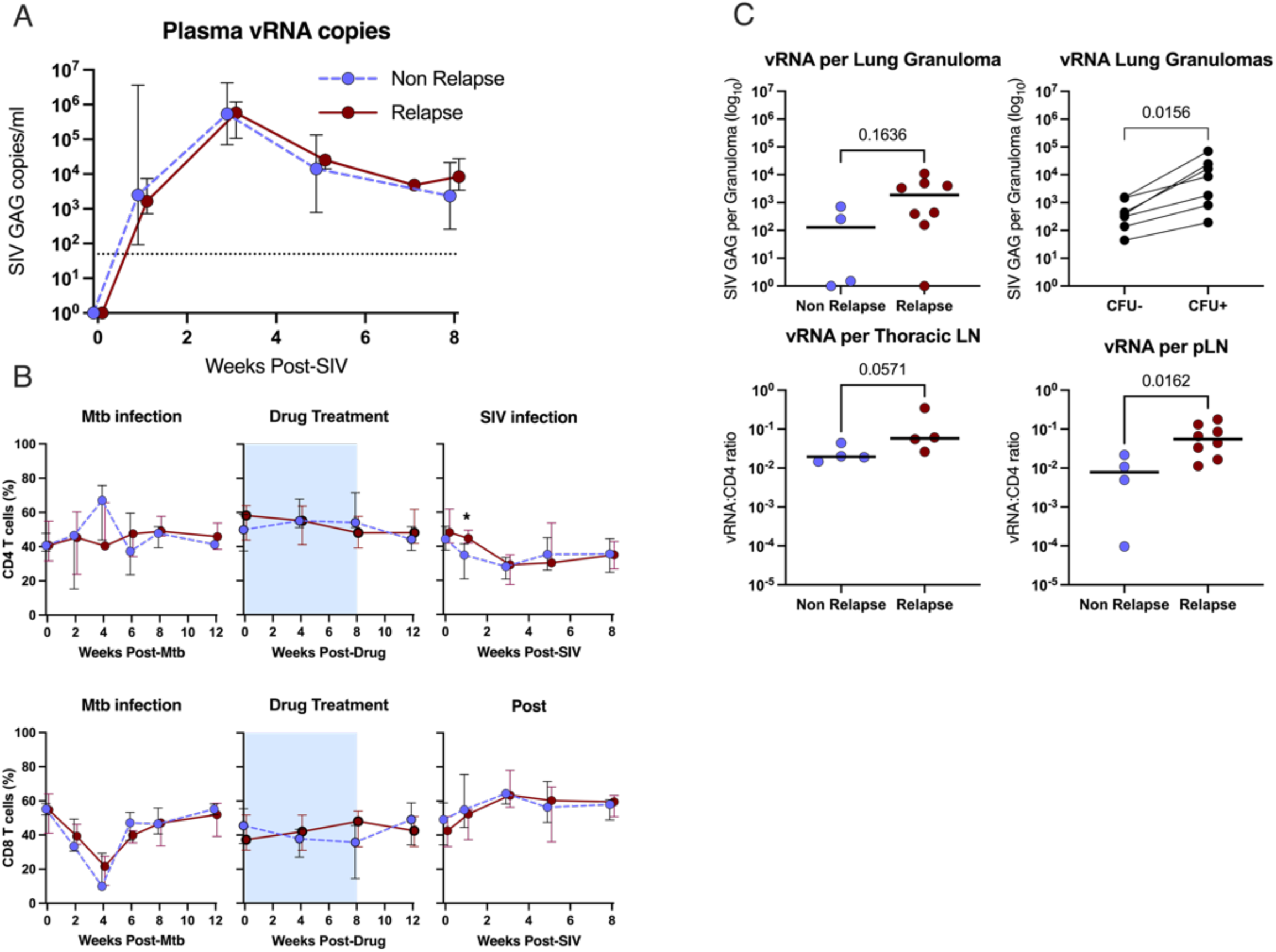
Peripheral plasma RNA levels and CD4 T cells frequencies do not correlate with relapse. (A) SIV plasma RNA copies/ml are shown over time among relapse and non-relapse animals during SIV infection. (B) Frequency of peripheral blood CD4 and CD8 T cells within the CD3 gate during Mtb infection (left column), drug treatment (blue background) with drug free period (middle columns), and during SIV infection (right column). Medians with IQR are shown. Mann-Whitney tests were run at each time point with no correction for multiple comparisons. p < 0.05: *. (C) The median SIV RNA levels in granulomas was similar between relapse and non-relapse animals. Greater median SIV RNA levels were observed from granulomas with viable Mtb (CFU+) compared to those without viable Mtb (CFU-). Greater SIV RNA:CD4 ratios are observed in relapse animals in both thoracic and peripheral lymph nodes (pLN). Top row: Each dot represents the median SIV/CD4 RNA ratio per animal. Bottom row: Each dot represents an individual lymph node. For unpaired group comparisons, p-value determined by Mann-Whitney. For paired data, p-value is determined by Wilcoxon matched-pairs signed rank test. A and B) 8 animals in relapse group, 4 animals in non-relapse group (not all animals represented at each time point).

**Supplemental Figure 9.**
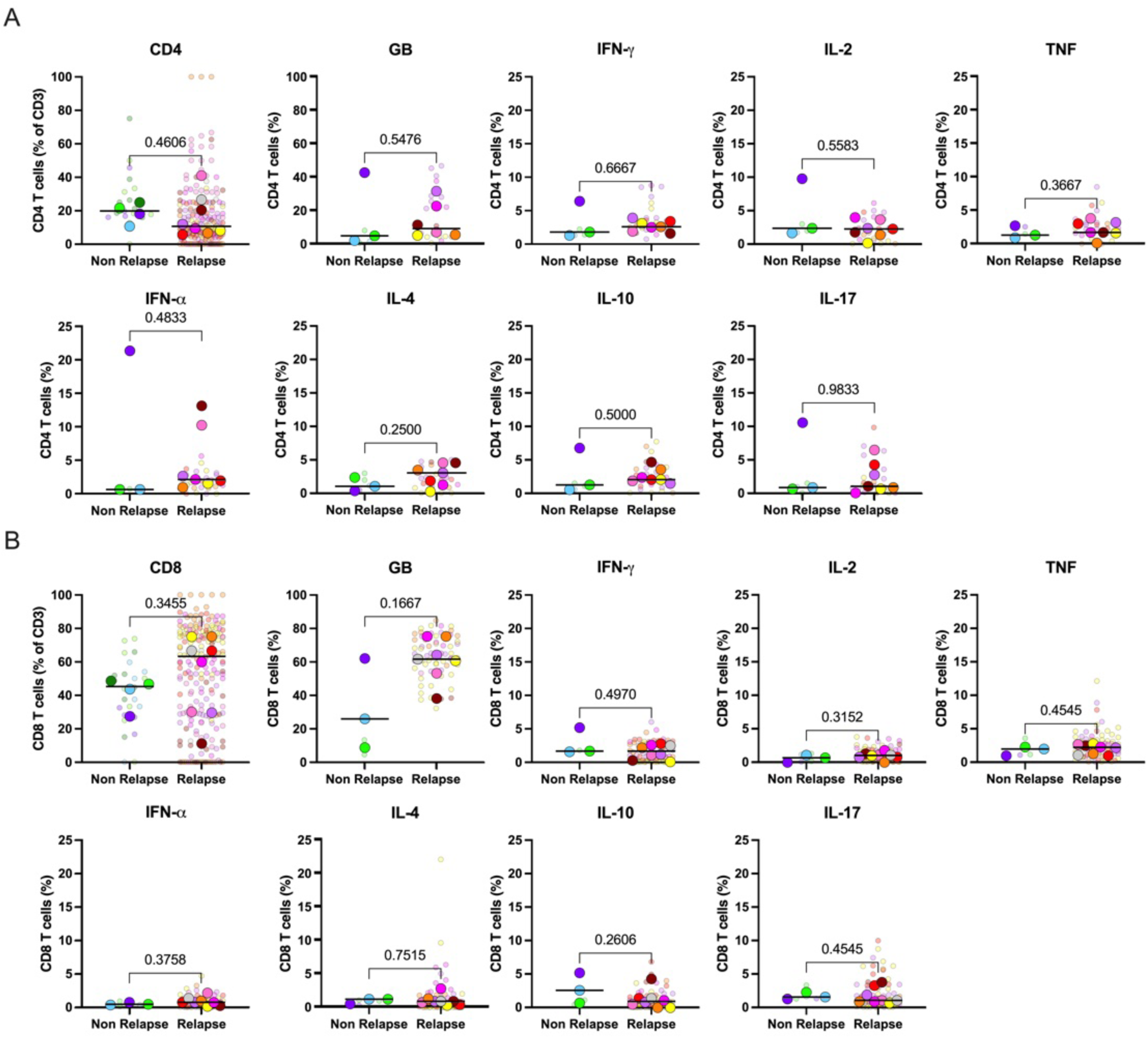
M**y**cobacterial **specific CD4 (A) and CD8 (B) T cell responses from granulomas are similar between relapse (n=3-4) and non-relapse animals (n=6-8).** Small, transparent circles represent individual granulomas; large circles reflect the median functional response from all granulomas analyzed per animal. Circles are colored by animal, and lines represent medians. P-values reflect Mann-Whitney analysis. (GB=granzyme B)

**Supplementary Figure 10.**
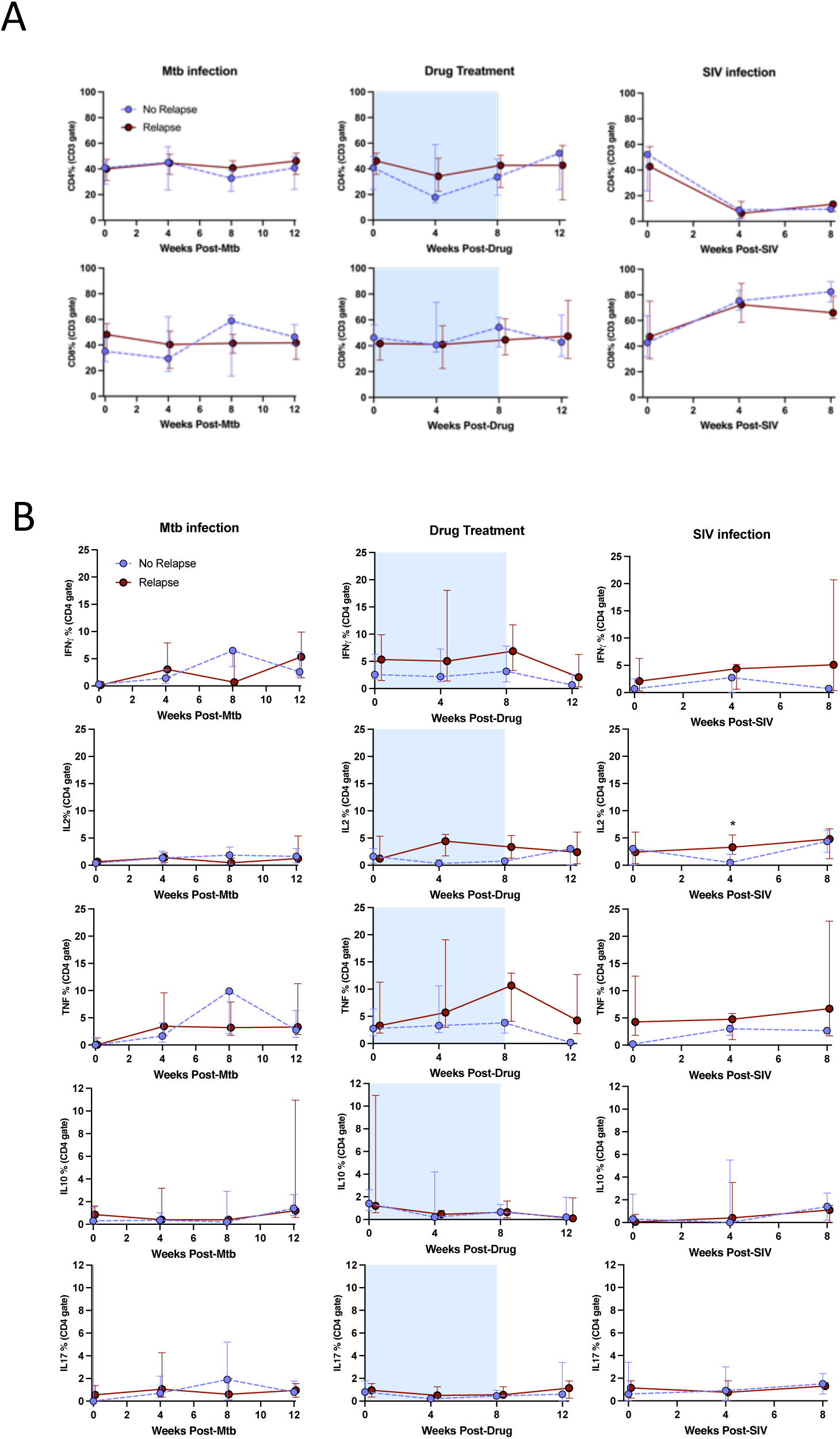
(A) Serial frequencies of CD4 and CD8 T cells in the airways. Serial CD4 (top) T cell frequencies and serial CD8 (bottom) T cell frequencies in the airway are shown during Mtb infection, drug treatment, and SIV infection among relapse (n=4) and non-relapse animals (n=8). (B) Serial frequencies of Th1 (IFN-³, IL-2, TNF), IL-10-, and IL-17-producing CD4 T cells in the airways during Mtb infection, drug treatment, and SIV infection among relapse (n=4) and non-relapse animals (n=8). Blue shaded area shows weeks of drug treatment, medians shown with IQR. Mann-Whitney tests were run at each time point with no correction for multiple comparisons. 0.05 < p < 0.10: #, p < 0.05: *.

**Supplementary Figure 11.**
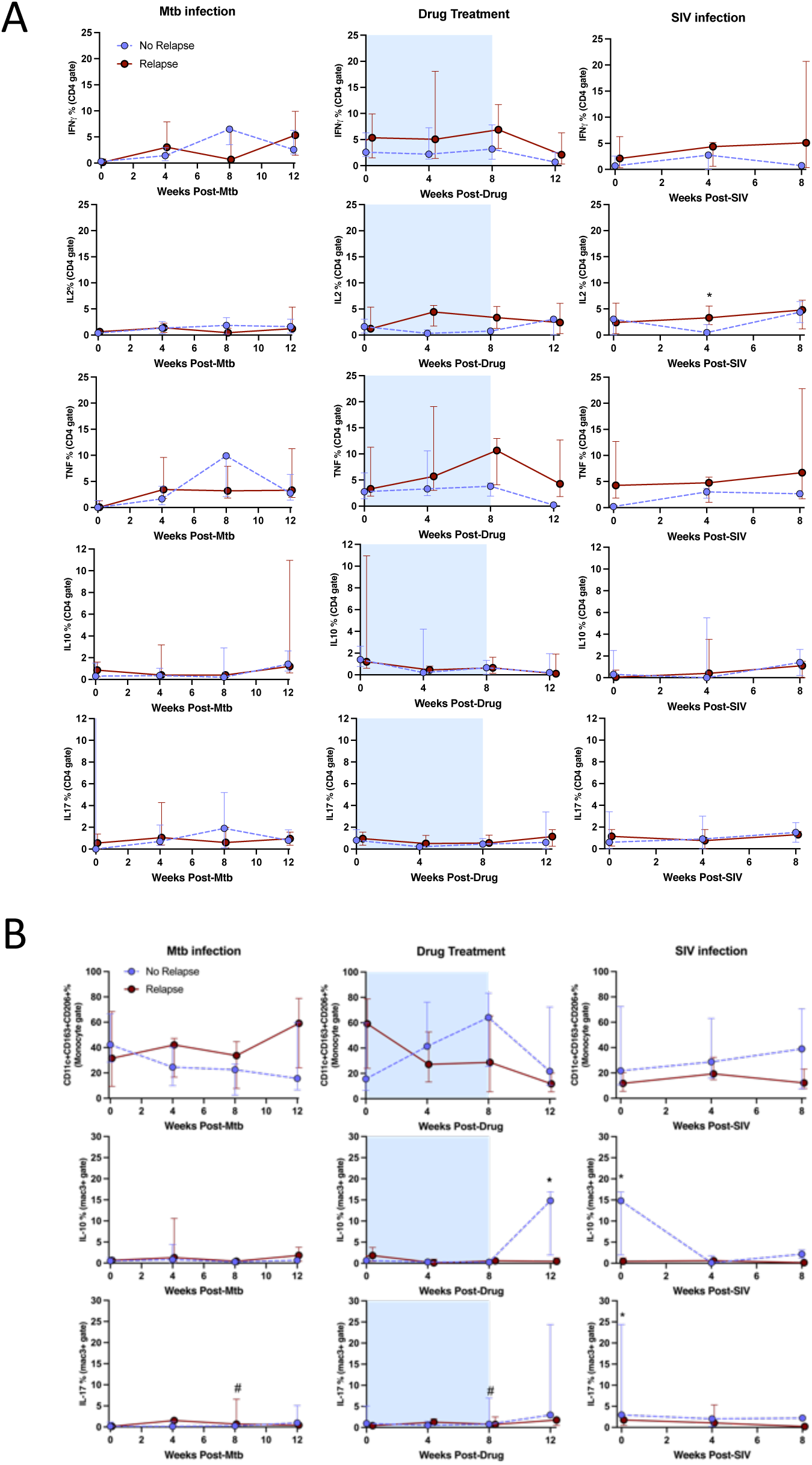
(A) Serial frequencies of Th1 (IFN-³, IL-2, TNF), IL10-, and IL17-producing CD8 T cells in the airways during Mtb infection, drug treatment, and SIV infection among relapse (n=4) and non-relapse animals (n=8). (B) Alveolar macrophage response in the airways over time. Top row: serial frequencies of alveolar macrophages (defined as CD11c+CD163+CD206+) in the airway. Middle row and bottom rows: Frequency of IL-10 and IL-17 producing alveolar macrophages over time. Blue shaded area shows weeks of drug treatment, medians shown with IQR Mann-Whitney tests were run at each time point with no correction for multiple comparisons. 0.05 < p < 0.10: #, p < 0.05: *.

**Supplementary Figure 12.**
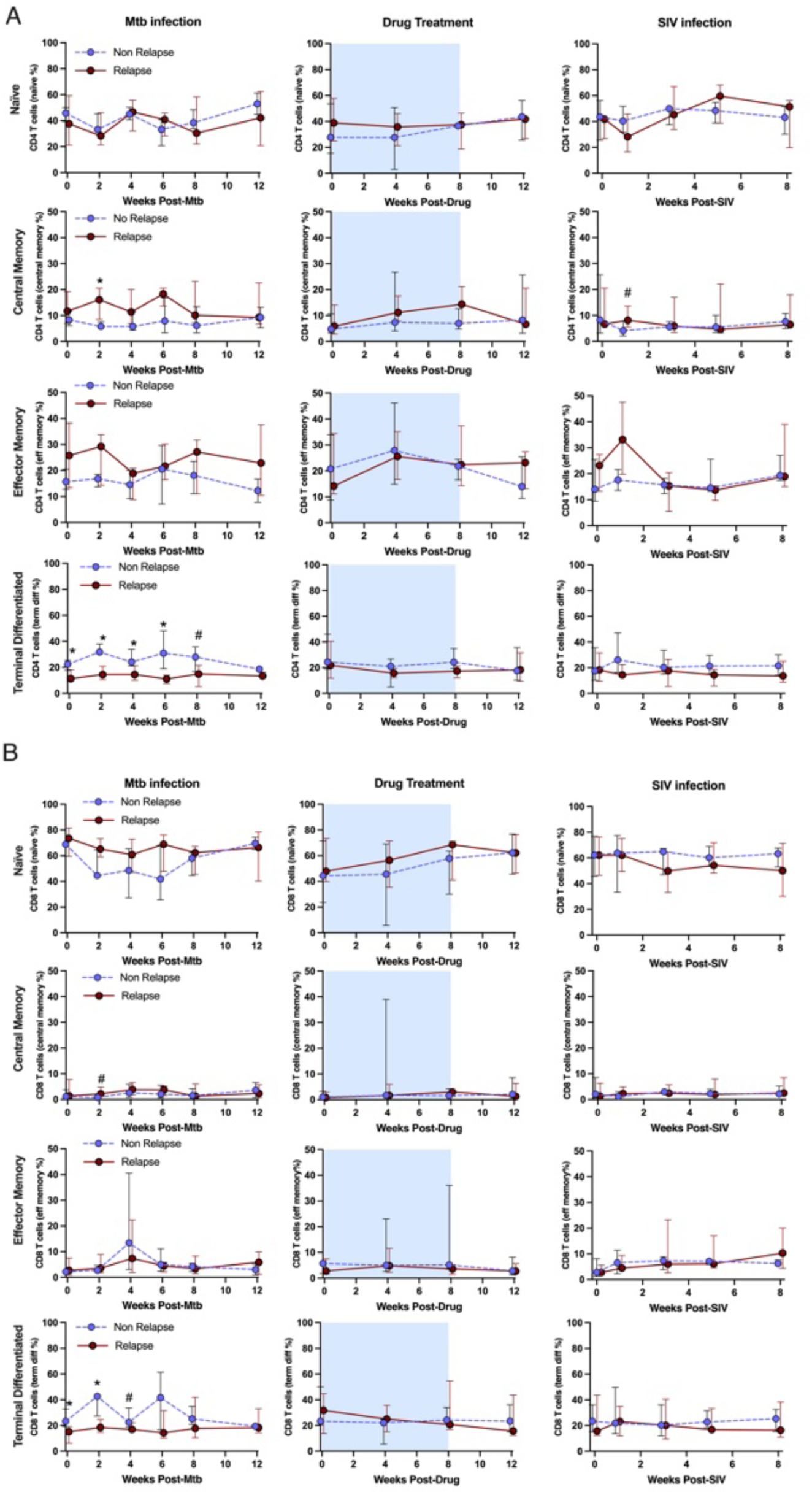
(A) Distribution of CD4 T cells in peripheral blood mononuclear cell by memory type during Mtb infection, drug treatment with drug free period, and SIV infection among relapse (n=8, n=7 for drug treatment graphs) and non-relapse (n=4) animals. Naïve (CD27+CD45Ra+), Central memory (CD27+CD45Ra-), Effector memory (CD27-CD45Ra-), Terminal differentiated (CD27-CD45Ra+). A lower frequency of terminal differentiated CD4 T cells appears during the first 6 weeks of Mtb infection among animals that would later develop relapse compared to those without relapse. (B) Distribution of CD8 T cells in peripheral blood mononuclear cell by memory type during Mtb infection, drug treatment with drug free period, and SIV infection among relapse (n=8) and non-relapse (n=4) animals. A lower frequency of terminal differentiated CD8 T cells appears during the first 2 weeks of Mtb infection among animals that would later develop relapse compared to those without relapse. Blue background in middle panels indicates drug treatment. Each dot reflects the median frequency and IQR shown. Mann-Whitney test used to compare groups at each time point. Unadjusted p-values: 0.05 < p < 0.10: #, p < 0.05: *.

**Supplementary Figure 13.**
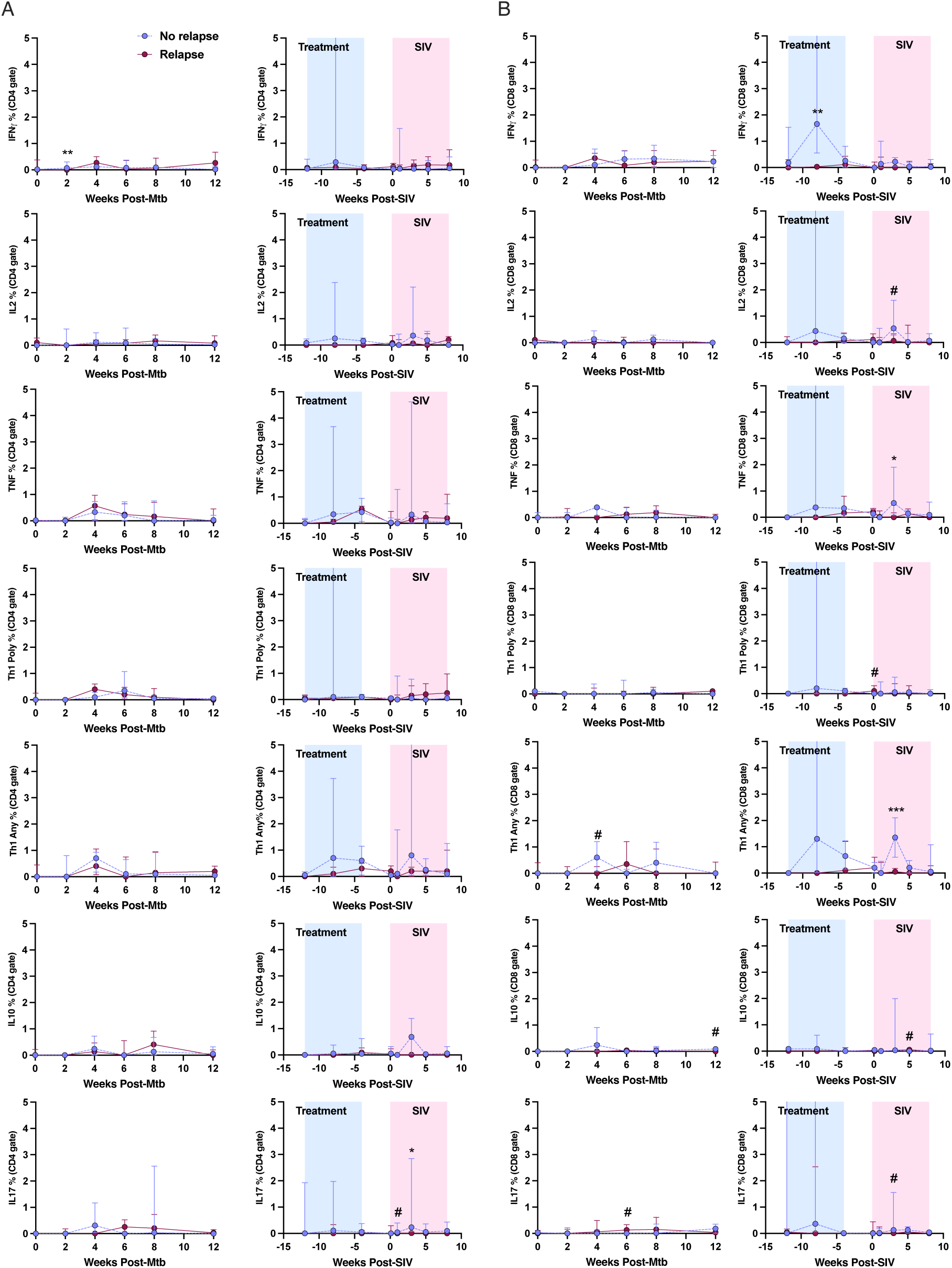
Serial frequencies of Mtb-specific, Th1 (IFN-³, IL-2, TNF), IL-10-, and IL-17-producing CD4 (A) and CD8 (B) T cells in the blood during Mtb infection, drug treatment, and SIV infection among non-relapse (n=4) and relapse (n=8) animals. Th1 Poly represents CD4 or CD8 T cells that produce two or more Th1 cytokines. Th1 Any represents CD4 or CD8 T cells that make at least one Th1 cytokine. Blue shaded area shows weeks of drug treatment, pink shaded area shows SIV-infection, medians shown with IQR. Mann-Whitney tests were run at each time point with no correction for multiple comparisons. 0.05 < p < 0.10: #, p < 0.05: *.

**Supplementary Figure 14.**
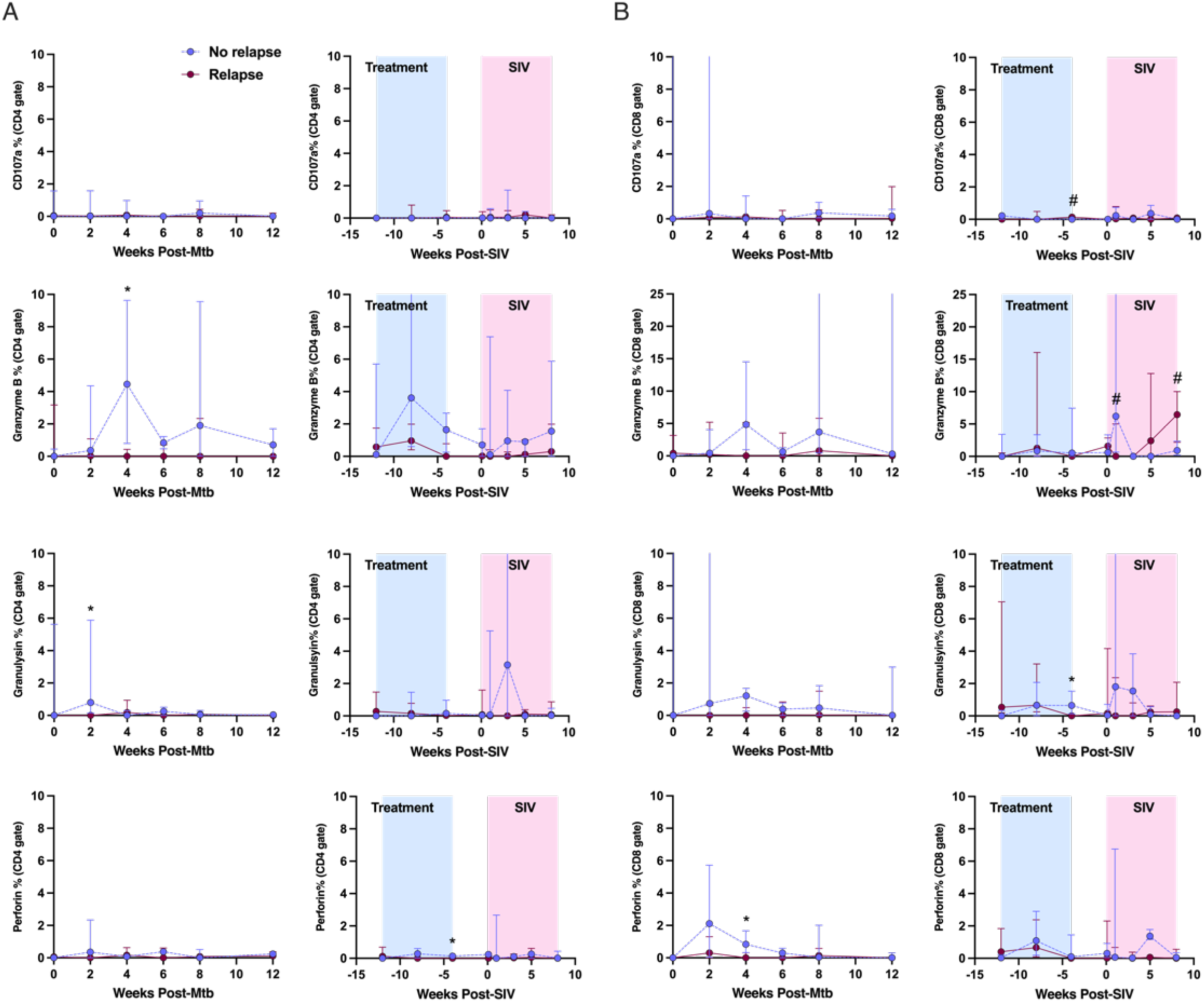
Serial frequencies of Mtb-specific CD4 (A) and CD8 (B) T cells with cytolytic characteristics in the blood during Mtb infection, drug treatment, and SIV infection among non-relapse (n=4) and relapse (n=8) animals. Blue shaded area shows weeks of drug treatment, pink shaded area shows SIV-infection, medians shown with IQR. Mann-Whitney tests were run at each time point with no correction for multiple comparisons. 0.05 < p < 0.10: #, p < 0.05: *.

**Supplementary Figure 15.**
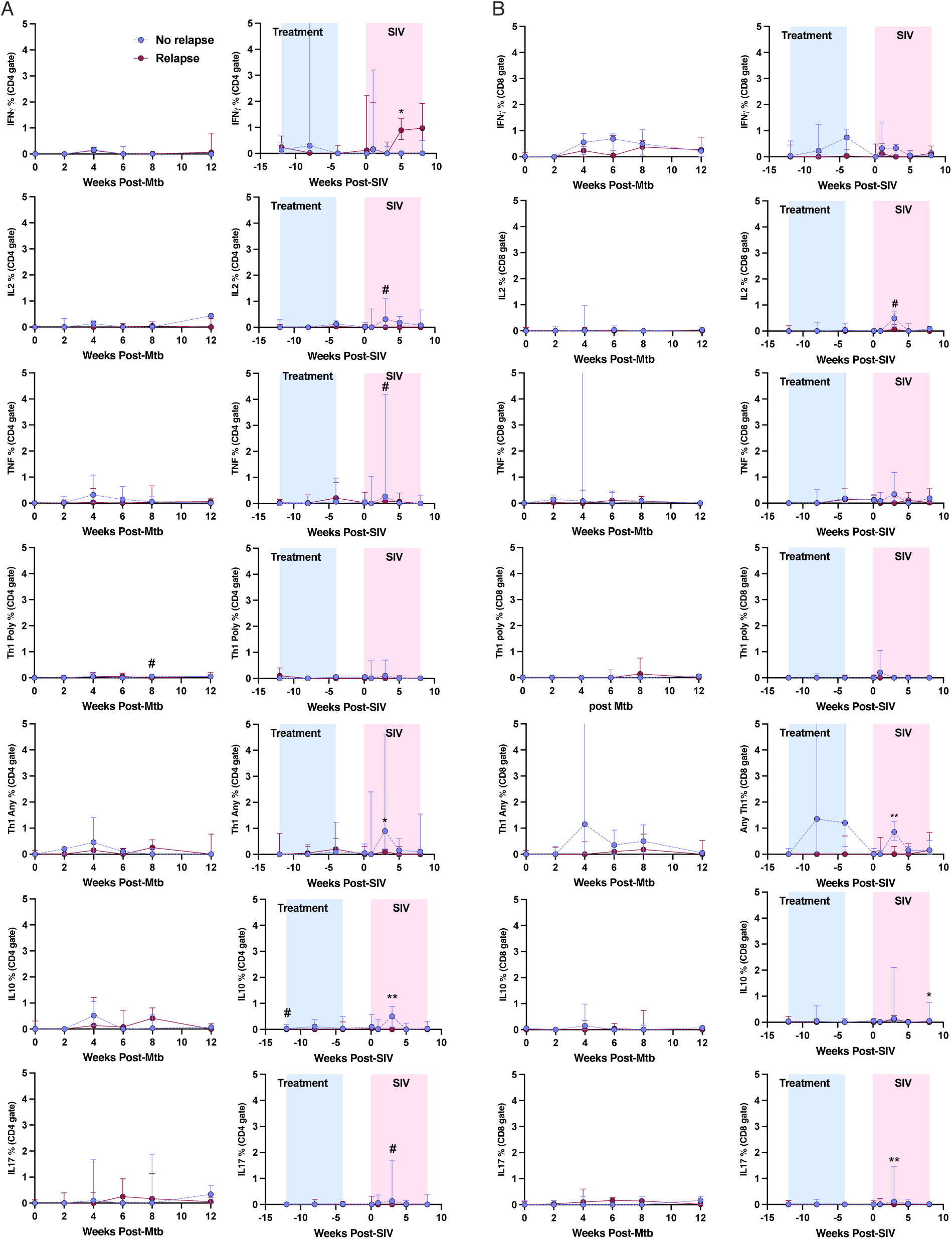
Serial frequencies of Mtb-specific, naïve (CD45Ra+CD27+), Th1-, IL10-, and IL17-producing CD4 (A) and CD8 (B) T cells in the blood during Mtb infection, drug treatment, and SIV infection among non-relapse (n=4) and relapse (n=8) animals. Th1 Poly represents CD4 or CD8 T cells that produce two or more Th1 cytokines. Th1 Any represents CD4 or CD8 T cells that make at least one Th1 cytokine. Blue shaded area shows weeks of drug treatment, pink shaded area shows SIV-infection. Mann-Whitney tests were run at each time point with no correction for multiple comparisons. 0.05 < p < 0.10: #, p < 0.05: *.

**Supplementary Figure 16.**
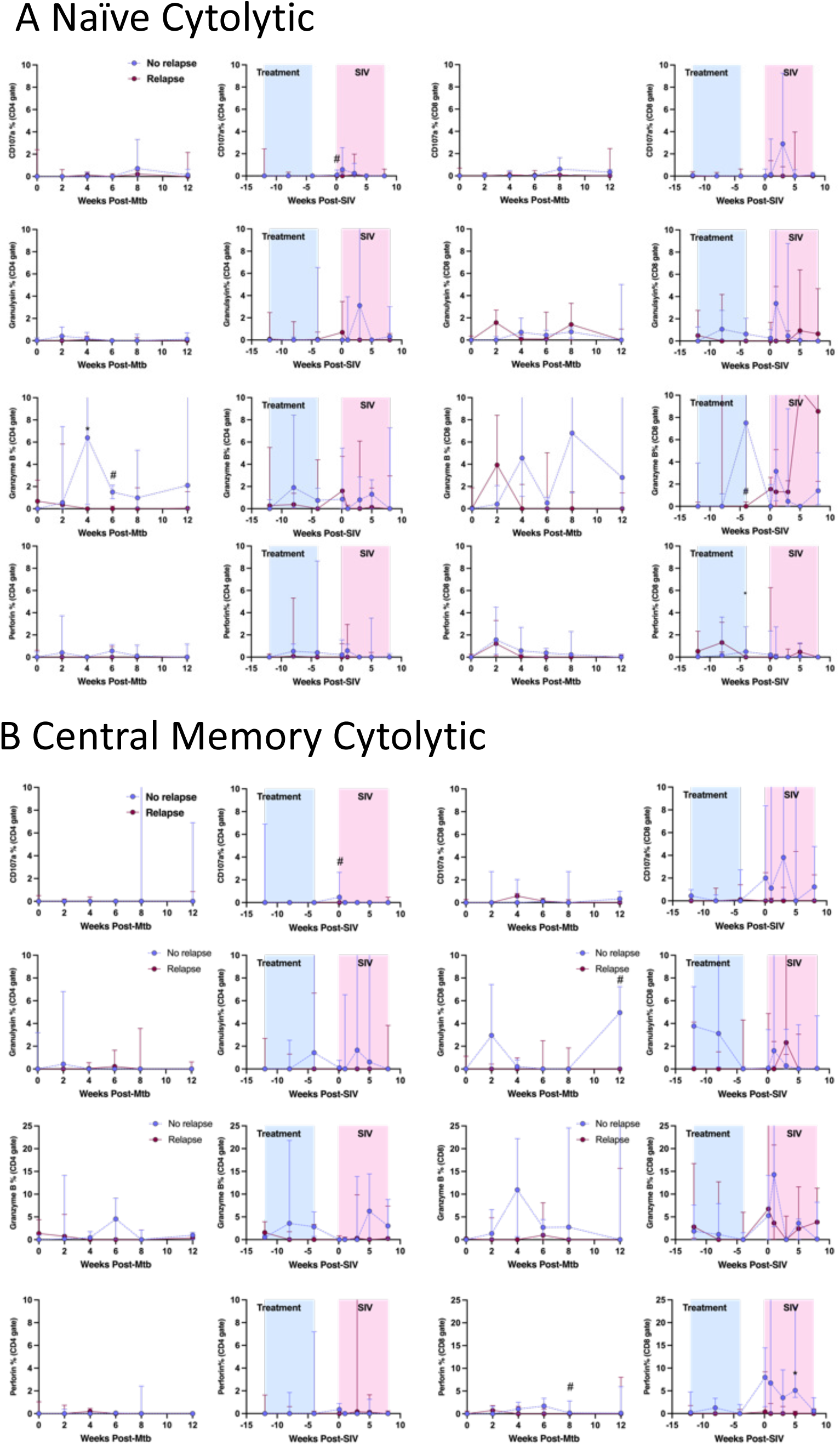
(A) Serial frequencies of Mtb-specific, naïve (CD45Ra+CD27) CD4 and CD8 T cells with cytolytic characteristics in the blood during Mtb infection, drug treatment, and SIV infection among non-relapse (n=4) and relapse (n=8) animals. (B) Serial frequencies of Mtb-specific, central memory (CD45Ra-CD27+) CD4 and CD8 T cells with cytolytic characteristics in the blood during Mtb infection, drug treatment, and SIV infection among non-relapse (n=4) and relapse (n=8) animals. Blue shaded area shows weeks of drug treatment, pink shaded area shows SIV-infection, medians shown with IQR. Mann-Whitney tests were run at each time point with no correction for multiple comparisons. 0.05 < p < 0.10: #, p < 0.05: *.

**Supplementary Figure 17.**
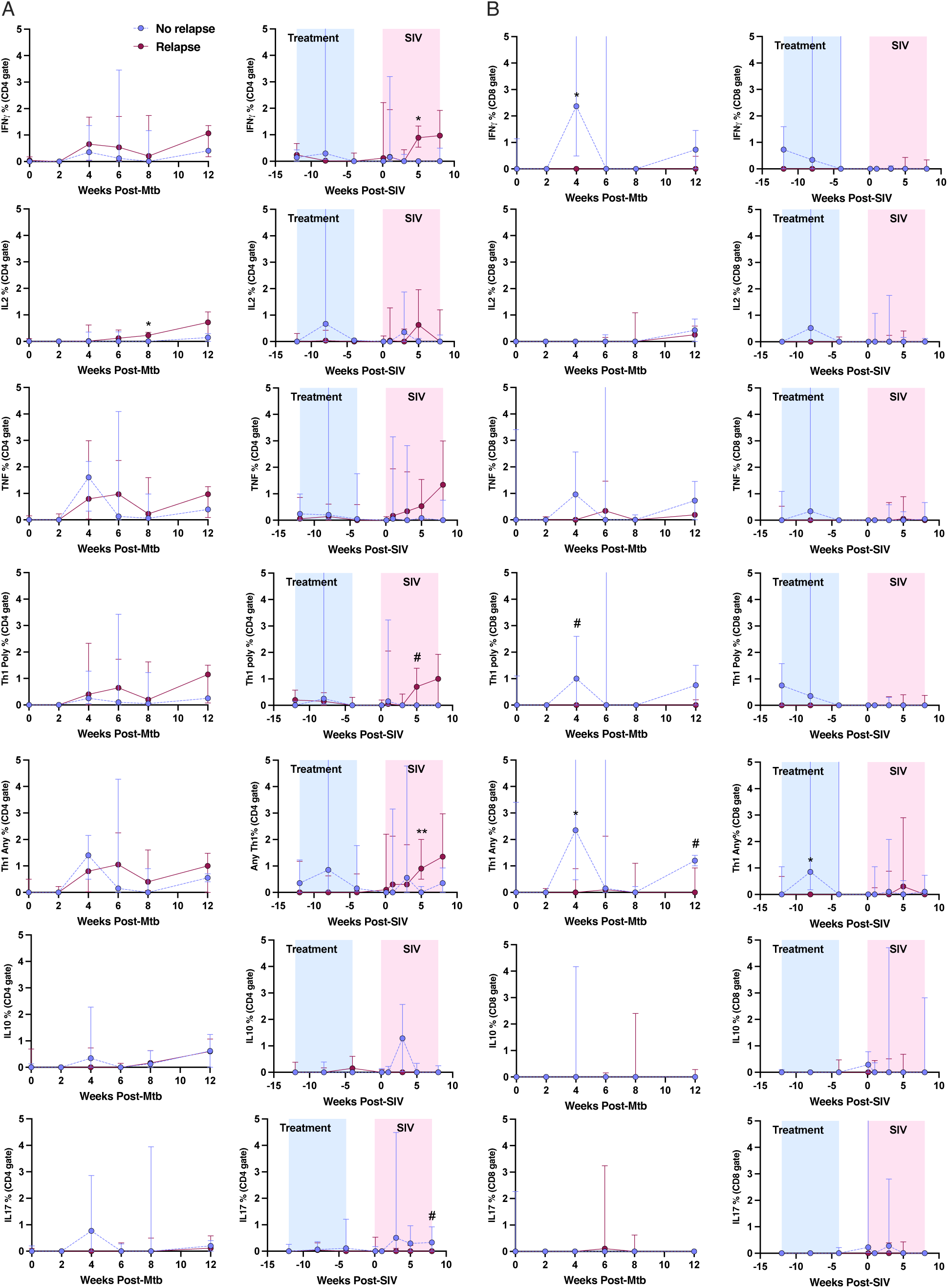
Serial frequencies of Mtb-specific, central memory (CD45Ra-CD27+), Th1-, IL10-, and IL17-producing CD4 (A) and CD8 (B) T cells in the blood during Mtb infection, drug treatment, and SIV infection among non-relapse (n=4) and relapse (n=8) animals. Th1 Poly represents CD4 or CD8 T cells that produce two or more Th1 cytokines. Th1 Any represents CD4 or CD8 T cells that make at least one Th1 cytokine. Blue shaded area shows weeks of drug treatment, pink shaded area shows SIV-infection, medians shown with IQR. Mann-Whitney tests were run at each time point with no correction for multiple comparisons. 0.05 < p < 0.10: #, p < 0.05: *.

**Supplementary Figure 18.**
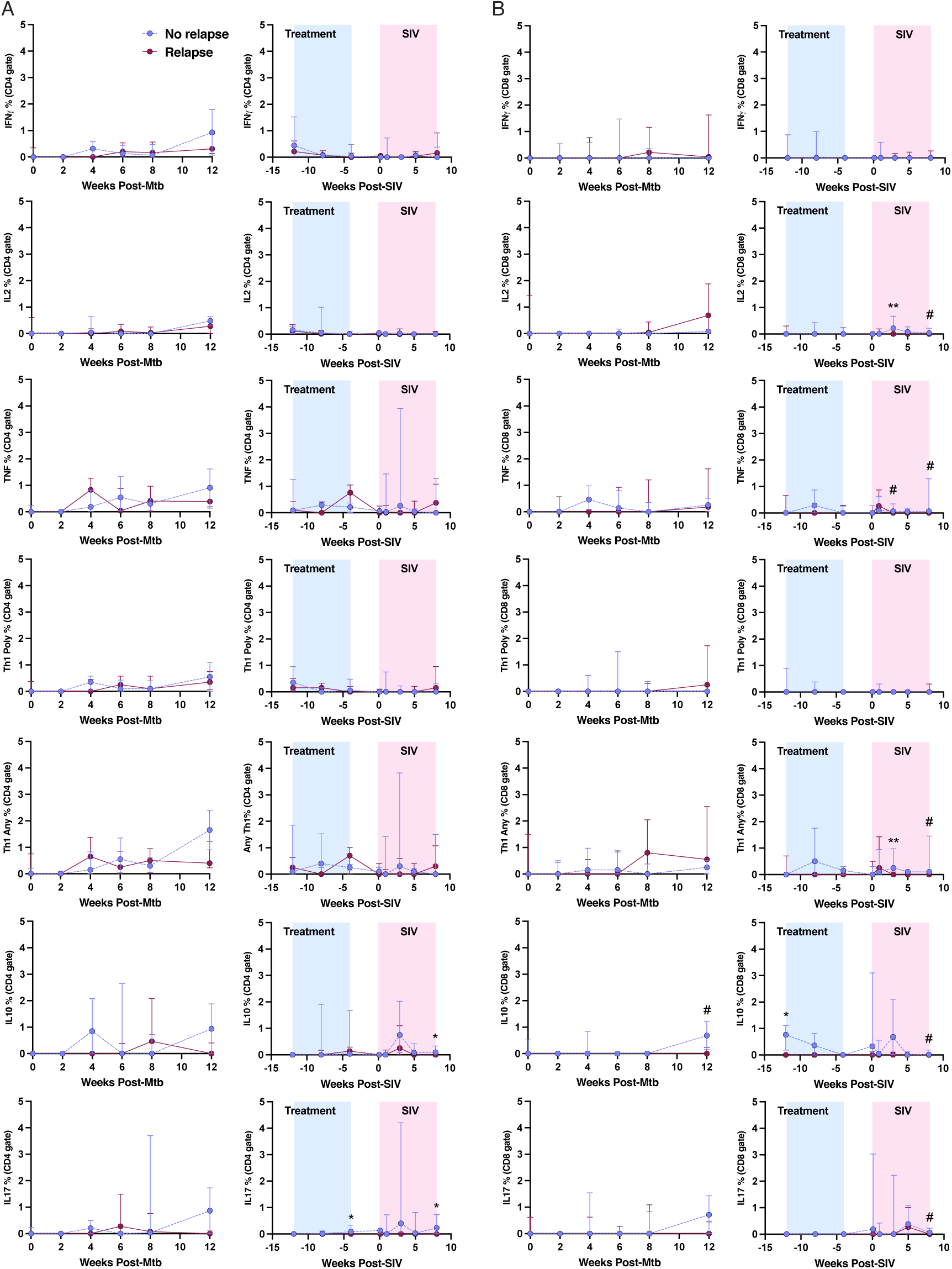
Serial frequencies of Mtb-specific, effector memory (CD45Ra-CD27-), Th1-, IL10-, and IL17-producing CD4 (A) and CD8 (B) T cells in the blood during Mtb infection, drug treatment, and SIV infection among non-relapse (n=4) and relapse (n=8) animals. Th1 Poly represents CD4 or CD8 T cells that produce two or more Th1 cytokines. Th1 Any represents CD4 or CD8 T cells that make at least one Th1 cytokine. Blue shaded area shows weeks of drug treatment, pink shaded area shows SIV-infection, medians shown with IQR. Mann-Whitney tests were run at each time point with no correction for multiple comparisons. 0.05 < p < 0.10: #, p < 0.05: *.

**Supplementary Figure 19.**
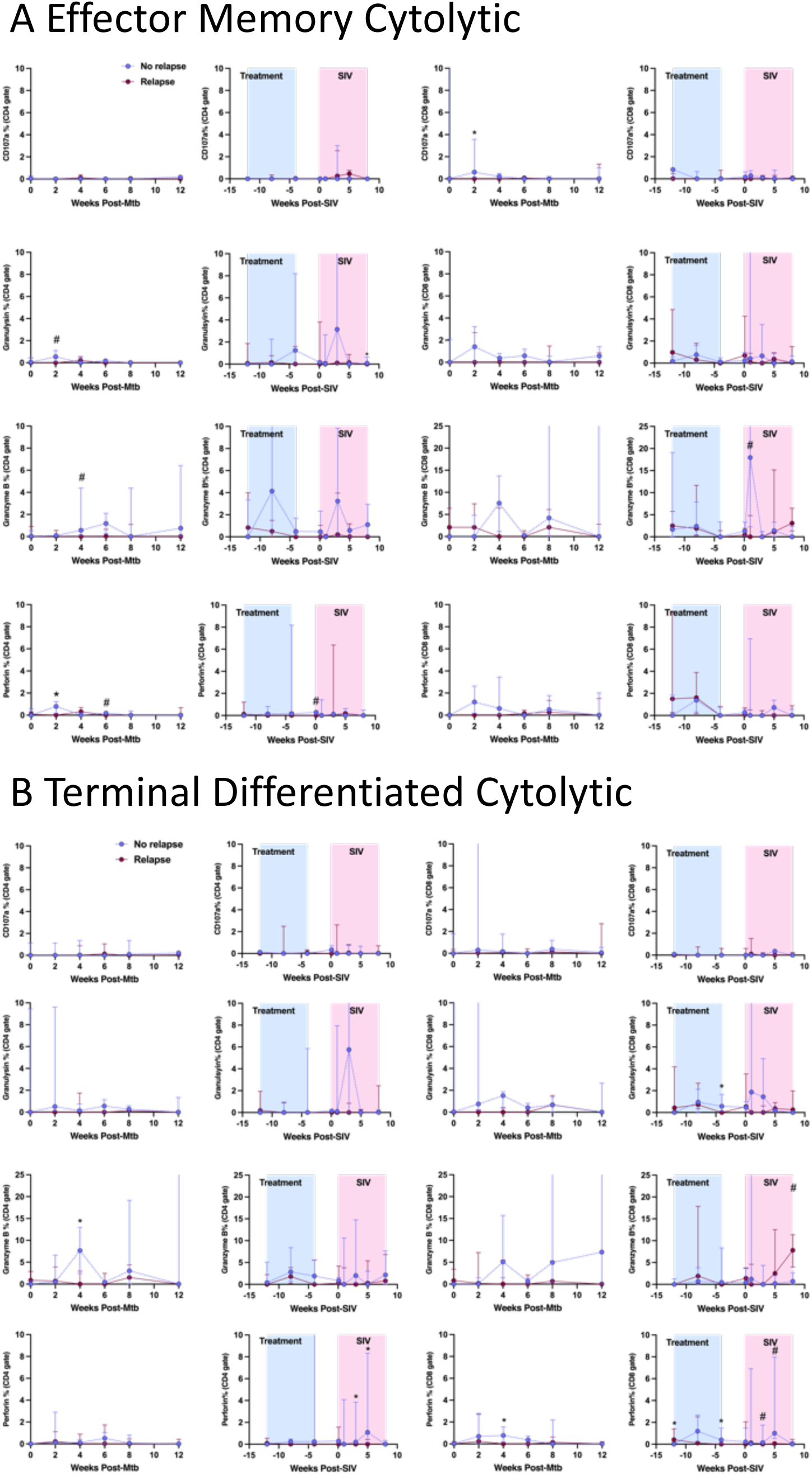
(A) Serial frequencies of Mtb-specific, effector memory (CD45Ra-CD27-) CD4 and CD8 T cells with cytolytic characteristics in the blood during Mtb infection, drug treatment, and SIV infection among non-relapse (n=4) and relapse (n=8) animals. (B) Serial frequencies of Mtb-specific, terminal differentiated (CD45Ra+CD27-) CD4 and CD8 T cells with cytolytic characteristics in the blood during Mtb infection, drug treatment, and SIV infection among non-relapse (n=4) and relapse (n=8) animals. Blue shaded area shows weeks of drug treatment, pink shaded area shows SIV-infection, medians shown with IQR. Mann-Whitney tests were run at each time point with no correction for multiple comparisons. 0.05 < p < 0.10: #, p < 0.05: *.

**Supplementary Figure 20.**
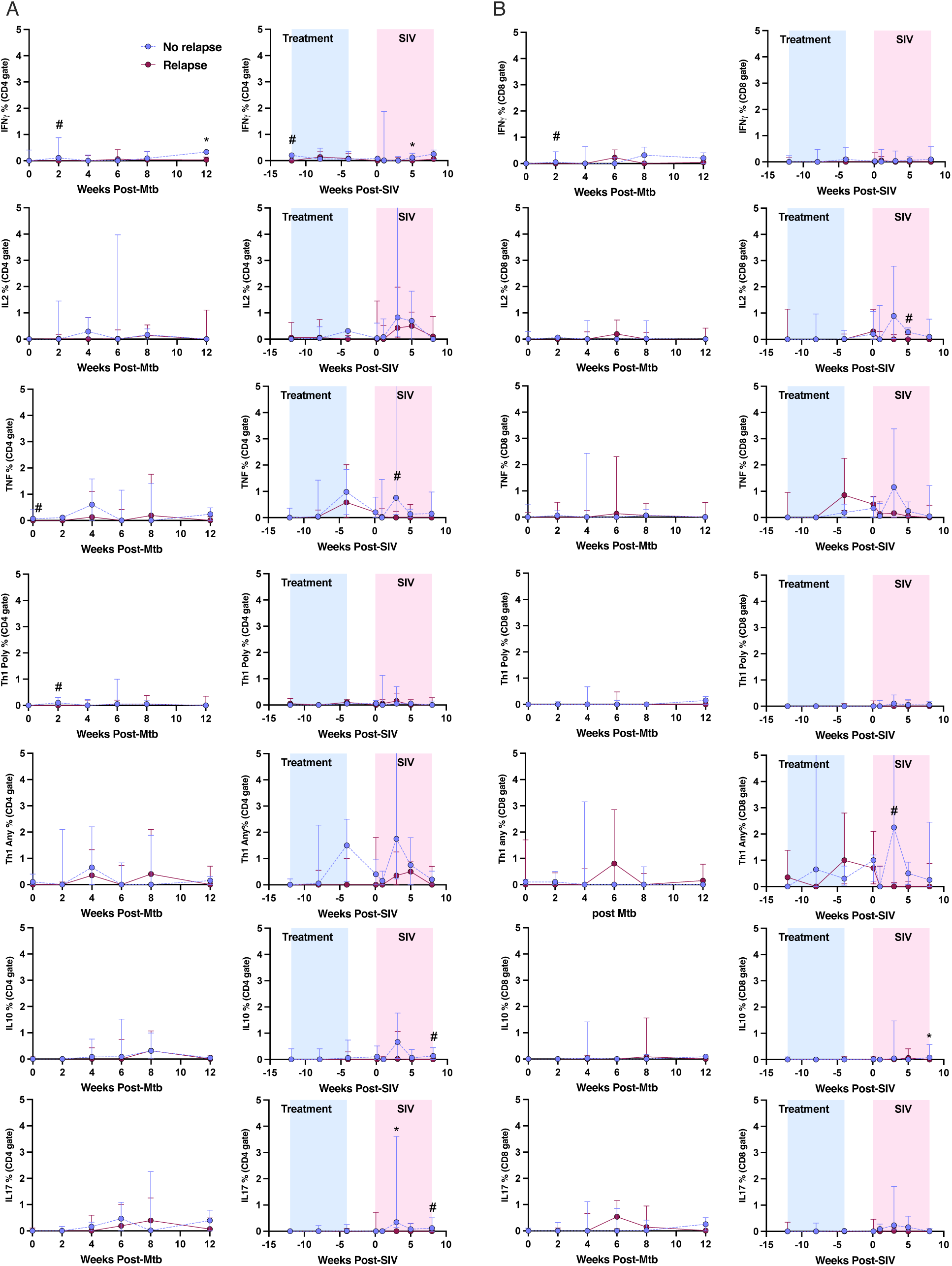
Serial frequencies of Mtb-specific, terminal differentiated (CD45Ra+CD27-), Th1-, IL10-, and IL17-producing CD4 (A) and CD8 (B) T cells in the blood during Mtb infection, drug treatment, and SIV infection among non-relapse (n=4) and relapse (n=8) animals. Th1 Poly represents CD4 or CD8 T cells that produce two or more Th1 cytokines. Th1 Any represents CD4 or CD8 T cells that make at least one Th1 cytokine. Blue shaded area shows weeks of drug treatment, pink shaded area shows SIV-infection, medians shown with IQR. Mann-Whitney tests were run at each time point with no correction for multiple comparisons. 0.05 < p < 0.10: #, p < 0.05: *.

**Supplementary Table 1.** Distribution of age and gender among the relapse and non-relapse animals.

|  | Total Animals | Age at Mtb Infection<br>(Years) |  | Gender<br>(Male) |
| --- | --- | --- | --- | --- |
|  | n | Mean | SD | Percent |
| **No Relapse** | 4 | 7.115753425 | 1.115 | 100.00% |
| **Relapse** | 8 | 6.980821918 | 0.723 | 87.50% |

**Supplementary Table 2.**
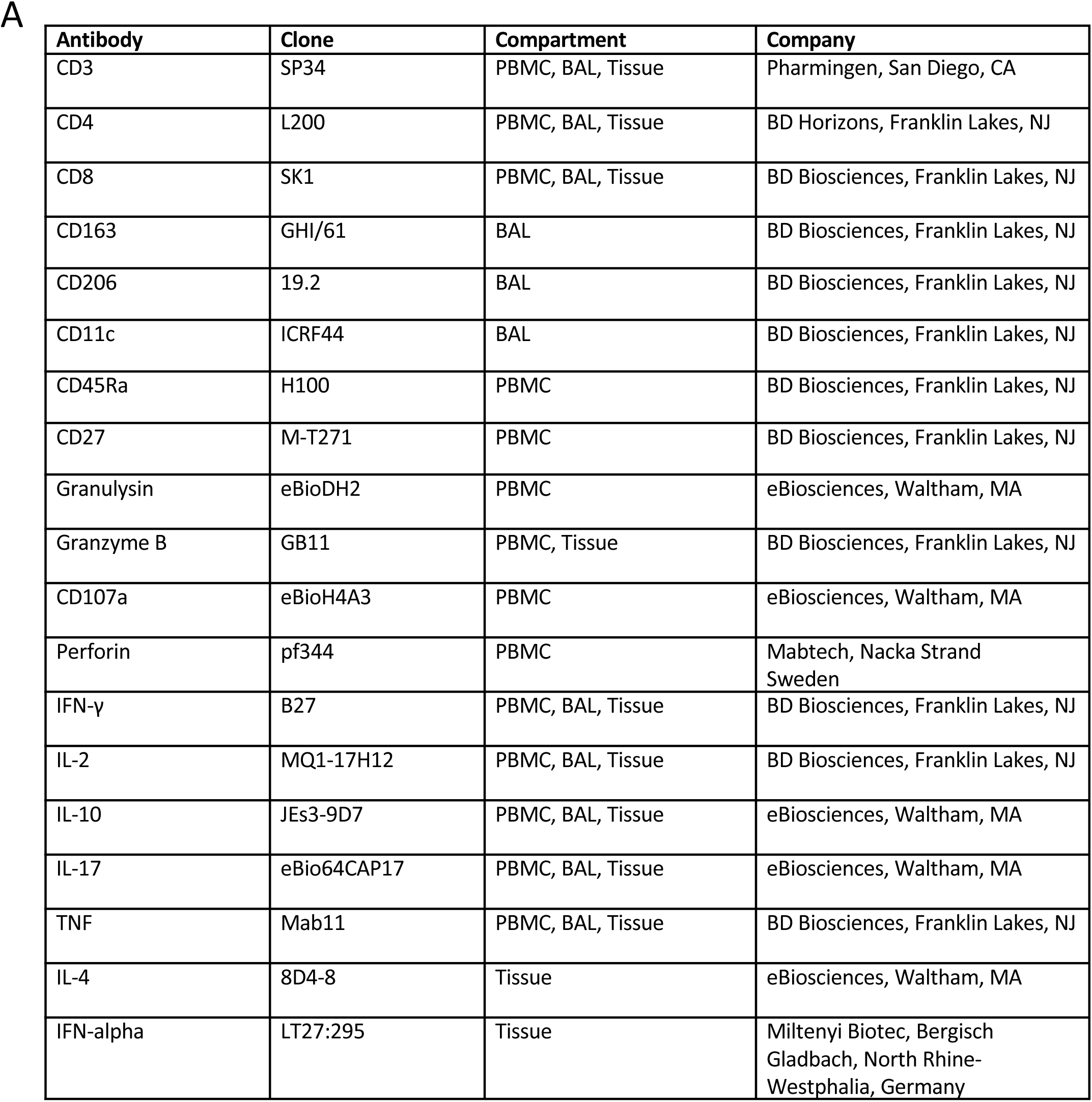
Antibodies used for flow cytometry assays.

| Antibody | Clone | Compartment | Company |
| --- | --- | --- | --- |
| CD3 | SP34 | PBMC, BAL, Tissue | Pharmingen, San Diego, CA |
| CD4 | L200 | PBMC, BAL, Tissue | BD Horizons, Franklin Lakes, NJ |
| CD8 | SK1 | PBMC, BAL, Tissue | BD Biosciences, Franklin Lakes, NJ |
| CD163 | GHI/61 | BAL | BD Biosciences, Franklin Lakes, NJ |
| CD206 | 19.2 | BAL | BD Biosciences, Franklin Lakes, NJ |
| CD11c | ICRF44 | BAL | BD Biosciences, Franklin Lakes, NJ |
| CD45Ra | H100 | PBMC | BD Biosciences, Franklin Lakes, NJ |
| CD27 | M-T271 | PBMC | BD Biosciences, Franklin Lakes, NJ |
| Granulysin | eBioDH2 | PBMC | eBiosciences, Waltham, MA |
| Granzyme B | GB11 | PBMC, Tissue | BD Biosciences, Franklin Lakes, NJ |
| CD107a | eBioH4A3 | PBMC | eBiosciences, Waltham, MA |
| Perforin | pf344 | PBMC | Mabtech, Nacka Strand<br>Sweden |
| IFN-γ | B27 | PBMC, BAL, Tissue | BD Biosciences, Franklin Lakes, NJ |
| IL-2 | MQ1-17H12 | PBMC, BAL, Tissue | BD Biosciences, Franklin Lakes, NJ |
| IL-10 | JEs3-9D7 | PBMC, BAL, Tissue | eBiosciences, Waltham, MA |
| IL-17 | eBio64CAP17 | PBMC, BAL, Tissue | eBiosciences, Waltham, MA |
| TNF | Mab11 | PBMC, BAL, Tissue | BD Biosciences, Franklin Lakes, NJ |
| IL-4 | 8D4-8 | Tissue | eBiosciences, Waltham, MA |
| IFN-alpha | LT27:295 | Tissue | Miltenyi Biotec, Bergisch<br>Gladbach, North Rhine-Westphalia, Germany |

